# Nested spatial and temporal modeling of environmental conditions associated with genetic markers of *Vibrio parahaemolyticus* in Washington state Pacific oysters

**DOI:** 10.1101/2022.01.03.471994

**Authors:** Brendan Fries, Benjamin J. K. Davis, Anne E. Corrigan, Angelo DePaola, Frank C. Curriero

## Abstract

The Pacific Northwest (PNW) is one of the largest commercial harvesting areas for Pacific oysters (*Crassostrea gigas*) in the United States. *Vibrio parahaemolyticus*, a bacterium naturally present in estuarine waters, accumulates in shellfish and is a major cause of seafood-borne illness. Growers, consumers, and public-health officials have raised concerns about rising vibriosis cases in the region. *V. parahaemolyticus* genetic markers (*tlh, tdh, trh*) were estimated using an MPN-PCR technique in Washington State Pacific oysters regularly sampled between May and October from 2005 to 2019 (N=2,836); environmental conditions were also measured at each sampling event. Multilevel mixed-effects regression models were used to assess relationships between environmental measures and genetic markers as well as genetic marker ratios (*trh:tlh, tdh:tlh*, and *tdh:trh*), accounting for variation across space and time. Spatial and temporal dependence were also accounted for in the model structure. Model fit improved when including environmental measures from previous weeks (1-week lag for air temperature, 3-week lag for salinity). Positive associations were found between *tlh* and surface water temp, specifically between 15°C and 26°C, and between *trh* and surface water temperature up to 26°C. *tlh* and *trh* were negatively associated with 3-week lagged salinity in the most saline waters (> 27 ppt). There was also a positive relationship between tissue temperature and *tdh*, but only above 20°C. The *tdh*:*tlh* ratio displayed analogous inverted non-linear relationships as *tlh*. The non-linear associations found between the genetic targets and environmental measures demonstrate the complex habitat suitability of *V. parahaemolyticus*. Additional associations with both spatial and temporal variables also suggest there are influential unmeasured environmental conditions that could further explain bacterium variability. Overall, these findings confirm previous ecological risk factors for vibriosis in Washington State, while also identifying new associations between lagged temporal effects and pathogenic markers of *V. parahaemolyticus*.

## Introduction

*Vibrio parahaemolyticus* is a gram-negative, halophilic bacterium naturally present in coastal waters around the world (CDC, 1998; Ansaruzzaman et al., 2005; Su and Liu, 2007; Kirs et al., 2011; Velazquez-Roman et al., 2014; Wu et al., 2014; Martinez-Urtaza et al., 2016). *V. parahaemolyticus* grows in oysters and accumulates when oysters are inactive. When oysters are active, they release more of the bacteria than they accumulate during filter feeding (FAO & WHO, 2021). Although *V. parahaemolyticus* is a thermolabile bacterium, oysters are most commonly consumed raw in the U.S. and so are the most common cause of seafood-borne bacterial gastroenteritis (Iwamoto et. al., 2010; Scallan et al., 2011) and are associated with the majority (90%) of *V. parahaemolyticus* infections (i.e., vibriosis) in the country (Froelich and Noble, 2016). Whereas symptoms are often mild and self-limiting, vibriosis can at times lead to life-threatening septicemia. The first vibriosis outbreak in the United States (U.S.) happened in 1971 and outbreaks have been consistently reported around the country since (Su and Liu, 2007).

In the U.S., the Pacific Northwest (PNW) region is known for its relatively high abundance of pathogenic *V. parahaemolyticus* strains (Paranjpye et al., 2012). The large tidal ranges in the PNW along with the use of intertidal harvesting practices result in oysters intended for consumption that have been exposed to air and direct sunlight for prolonged time-periods (Jones et al., 2016). The first large vibriosis outbreak in the PNW occurred in 1997; 209 people became ill from eating shellfish harvested in the PNW and one person died (CDC, 1998). There have been additional outbreaks in the PNW since that time along with an increased incidence of intermittent cases in Washington State where the majority of shellfish are harvested in the PNW (USFDA, 2005; National Marine Fisheries Service et al., 2017; Washington Disease Control and Health Statistics, 2018). In spite of various efforts by the Washington State Department of Health (WDOH) and shellfish growers to limit oyster harvest by time and place and to employ strict post-harvest controls, instances of vibriosis have continued to rise over time, (Paranjpye et al., 2012; Washington State Board of Health, 2015). Cases in similar estuarial environments in other parts of the world now occur both earlier and later in the year (Muhling et al., 2017; Vezzulli et al., 2013), and thus will likely also start to occur in the PNW. Climatic changes, especially those at higher latitudes, may be responsible for these trends, possibly from the increase in sea temperatures, given the well-reported positive correlation between *V. parahaemolyticus* abundance and water temperature (Baker-Austin et al., 2017; Sterk et al., 2015; Vezzulli et al., 2016).

The bacterium population found in PNW oysters is comprised of a wide array of strains that partake in consistent genetic recombination (Hazen et al., 2010; Paranjpye et al., 2012; Turner et al., 2013). Not all *V. parahaemolyticus* strains can cause vibriosis, although all *V. parahaemolyticus* bacteria have a specific thermolabile hemolysin gene (*tlh*). Frequent markers of pathogenicity found in *V. parahaemolyticus* bacteria are the thermostable direct hemolysin and the thermostable direct-related hemolysin genes (*tdh* and *trh*) (Shirai et al., 1990; Panicker et al., 2004). The bacterium strains often found in PNW waters, predominantly those containing the *tdh* and *trh* genetic markers, are regularly connected with vibriosis cases in Washington (Martinez-Urtaza et al., 2013; Paranjpye et al., 2012; Turner et al., 013; Velazquez-Roman et al., 2014; Xu et al., 2017, Davis et al., 2020). Although neither of the genes always predict or are required for pathogenicity, they have been frequently associated with type III secretion systems that can lead to human infection (Paranjpye et al., 2012; Whistler et. al., 2015). For this reason, they are often used to gauge the measure of virulence factors in environmental isolates (Zhang and Orth, 2013).

An array of environmental conditions have previously been associated with the presence and abundance of *V. parahaemolyticus* in estuarial environments (DePaola et al., 2003; Turner et al., 2014; Johnson, 2015; Paranjpye et al., 2015; Davis et al., 2017). For example, water temperature is used as the primary environmental input in the U.S Food and Drug Administration’s risk assessment for *V. parahaemolyticus* in oysters (USFDA, 2005). While the bacterium can only survive in saline waters, its concentrations at higher salinity levels vary widely due at least in part to the other (a)biotic conditions of the water column, including variability in oyster activity. It is difficult, however, to identify broadly applicable relationships between environmental determinants and abundance of the bacterium as many of these associations vary significantly across geographic regions, time periods, and by the genetic marker of interest (DePaola et al., 2003; USFDA, 2005; Turner et al., 2013; Johnson, 2015; Paranjpye et al., 2015; Davis et al., 2017). Previous findings in the PNW have laid the groundwork to characterize space–time residual dependencies for the relationships between environmental measures and *V. parahaemolyticus* abundance in the region (Flynn et al., 2019; Paranjpye et al., 2015). Statistical assessments accounting for spatial heterogeneity and autocorrelation is also important to consider across the varied aquatic “zones” found in Washington state (Davis et al., 2020).

Given the vibriosis health concerns in Washington, the economic impact on some of the largest oyster harvesters in the U.S. and expected increases in illness rates as higher latitude waters continue to warm (Baker-Austin et al., 2017), further inquiry into the environmental determinants of *Vibrio* bacterium in the region is merited. This study utilizes a large dataset of regularly monitored harvesting sites and public shellfish collection beaches in the state of Washington. The dataset used in this study contains one of the most comprehensive assortments of oyster samples ever analyzed for *V. parahaemolyticus* and spans a considerable range of locations and years. The current dataset, therefore, allows for an in-depth examination of the independent associations between genetic markers of the bacterium and aquatic environmental measurements. This study specifically aimed to evaluate the relationships between absolute and relative abundance of *V. parahaemolyticus* strains carrying selected pathogenic genetic markers with temperature measures and salinity, while accounting for temporal autocorrelation, in Washington State.

## Methods

### Oyster Sampling and Processing

Starting in 2005, the Washington Department of Health (WDOH) has regularly sampled Pacific oysters (*Crassostrea gigas*) in order to quantify the abundance of *V. parahaemolyticus* genetic markers. Intertidal shellfish harvesting beds were sampled each year, as frequently as once a week, between May and October from 2005 to 2019. Sampling was performed primarily in Hood Canal and South Puget Sound, with additional samples taken in the coastal bays and northern waters (comprised of Samish Bay and Drayton Harbor) (**Figure 1**).

**Fig. 1.**
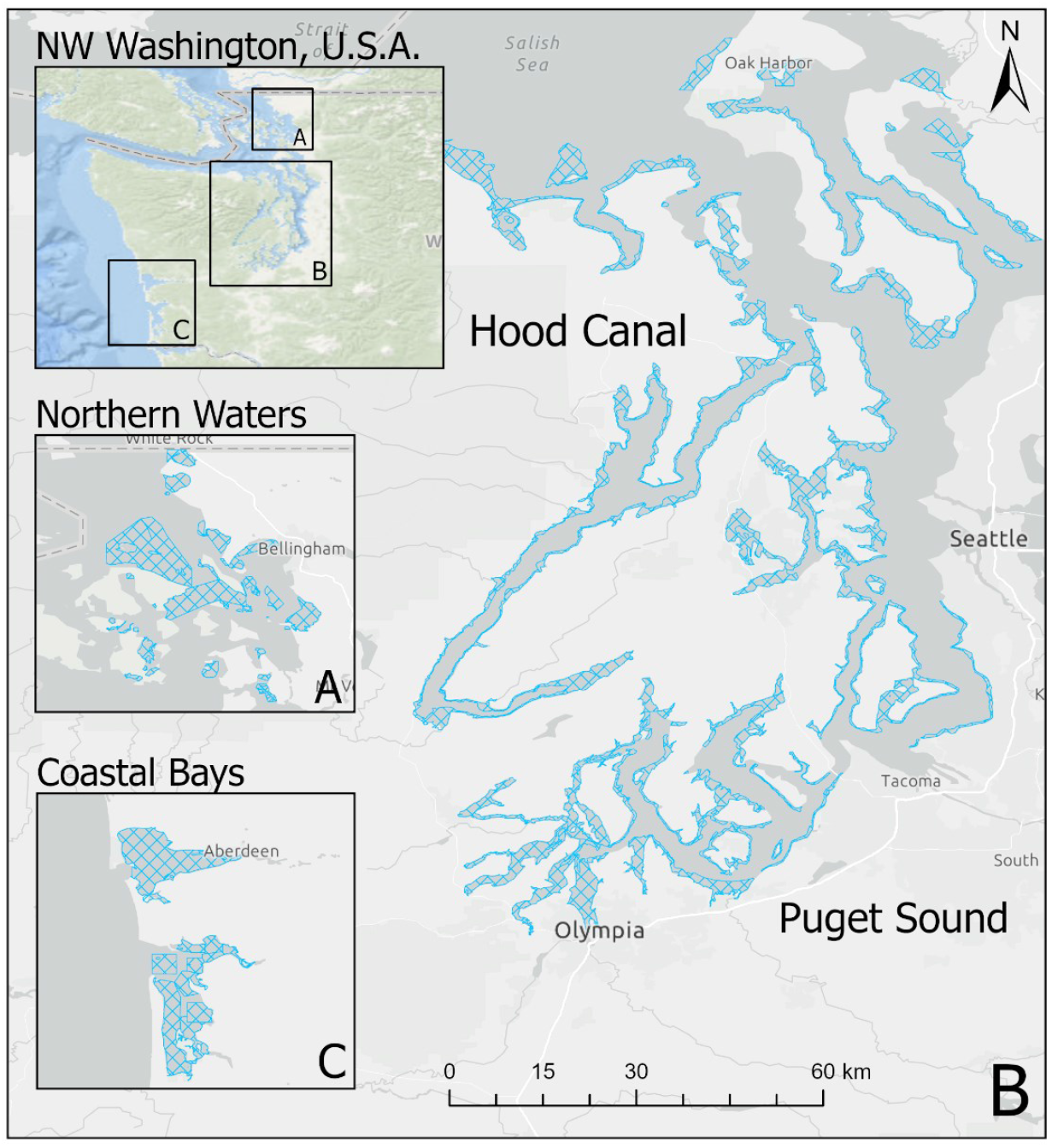
Oyster growing areas by zone in Washington state. Figure shows growing areas (blue hatched polygons) in (A) Coastal Bays, (B) Hood Canal and South Puget Sound, and (C) Northern Waters including Samish Bay. Puget Sound and Hood Canal represent separate oyster “zones” for the purpose of this study.

Each sample consisted of 13-18 live oysters approximately 10 cm in length. When feasible, oysters were collected during the receding tide (*i*.*e*., when no longer submerged). During collection, oysters were rinsed in seawater, sealed in a bag, and placed in a chilled insulated container kept at 2-8 °C. One additional oyster was shucked, and its tissue temperature was recorded before being discarded. At the time of collection, shoreline water, surface water (measured where depth exceeded 0.6 m), and ambient air temperature were recorded with a thermometer. Salinity from water samples was measured with a refractometer in parts per thousand (ppt). Oyster samples were processed within 24 hours of collection. Whole oyster tissue within a sample was homogenized and *V. parahaemolyticus* genetic markers enumerated using a most-probable-number per gram (MPN/g) based real-time PCR assay. Details of this assay have been described previously (Flynn et al., 2019; Glover II, 2015), and the MPN-PCR analytical results have been validated. Briefly, hemolysin genetic markers *tdh* and *tlh* were targeted across all sampling years, with *trh* also being targeted beginning in 2014. The analytical limit of detection for all three markers was 0.3 MPN/g with the upper limit of quantification being 110,000 MPN/g. Note that the assay was limited to separately assessing the gross abundance of each genetic marker; therefore, it was unable to distinguish between bacterium strains and/or sequence types.

The current work included samples collected between 2005-2019. Sampling in 2009 was unique in that a non-systematic sampling schema was used, and therefore samples from this year were excluded from the described analyses. After exclusion of 2009, our dataset resulted in 3,137 total oyster samples, entries with lab errors or data errors were excluded (9.5%), leaving 2,836 samples for the analysis. Sample sites were categorized based on the time and geographic location of sampling. Sample-Site-Year-Groups (SSYGs) were constructed based on sample sites in a specific year, for example the “Oyster Bay” sampling site was only sampled in yeas 2013, 2014, and 2016 and the exact coordinates of sampling shifted between 2014 and 2016. From this one sampling site, three SSYGs were created: Oyster Bay 2013, Oyster Bay 2014, and Oyster Bay 2016; with corresponding separate geocoordinates. Locations were nested by zone (*i*.*e*., South Puget Sound, Hood Canal, Coastal Bays, and the Northern Waters) and sampling site (**Figure 2**). Time as a variable was measured as weeks and years, with within-season successive weeks as linear time series nested in their respective years, such as 18 weekly entries in the Hood Canal zone in 2005 form a portion of the entire longitudinal Hood Canal dataset. Exploratory analyses included scatterplots and boxplots of each environmental and genetic variable to compare univariate and bivariate relationships across the spatial and temporal variables. Non-linear trends were identified using non-parametric local regressions (LOESSs). For exploratory analyses, genetic variable assay values below the limit of detection were halved and those above the upper limit of quantification were doubled.

**Fig. 2.**
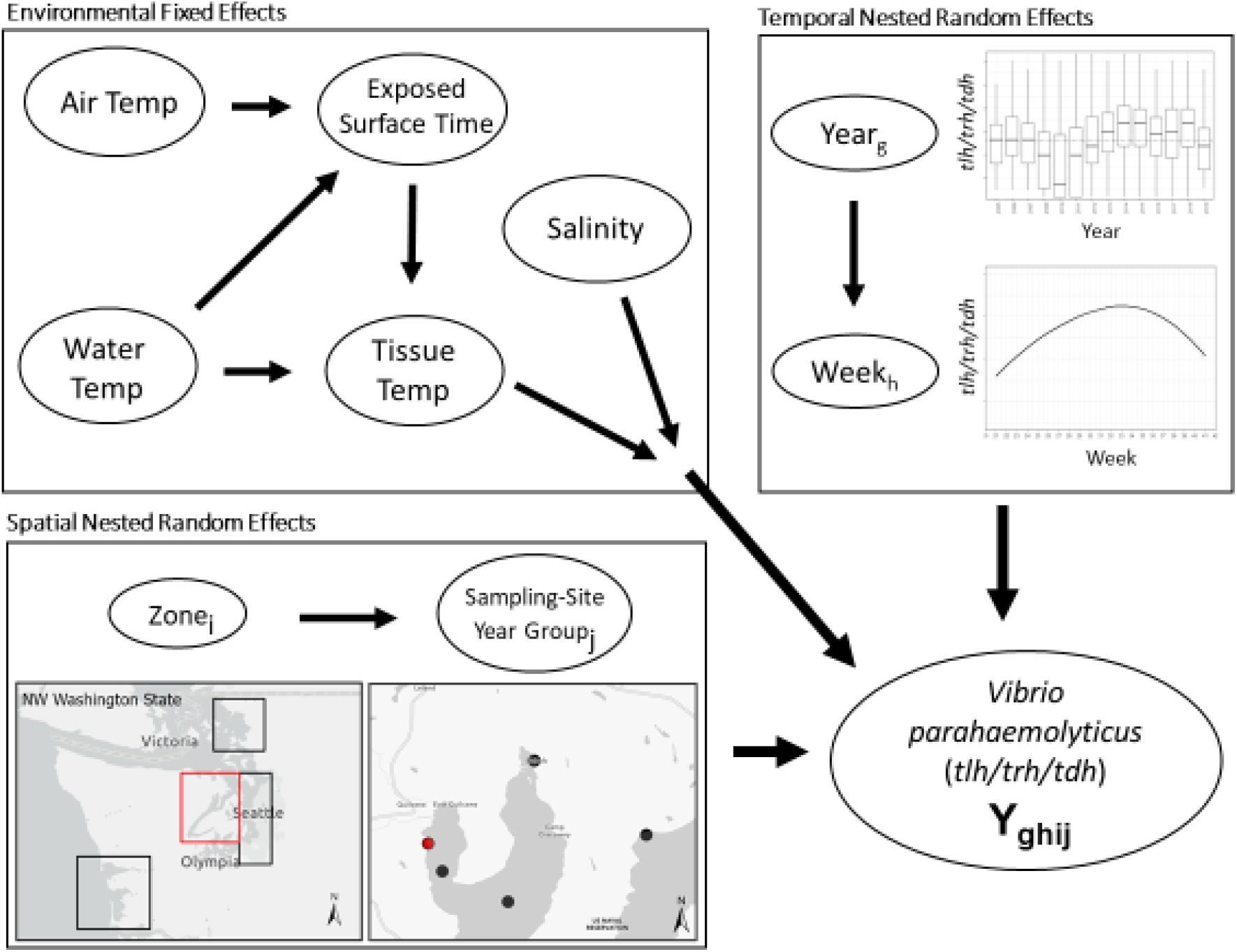
Schematic illustration of base nested model structure. First level environmental conditions fixed effects are modified by higher level spatial and temporal random effects. Spatial random effect of SSYG (*j*) is nested within effect of oyster growing zone (*i*) in Washington state. Temporal random effect of sampling week (*h*) is nested within random effect of year (*g*). Spatial and temporal model levels modify relationship between environmental conditions and concentrations of *Vibrio parahaemolyticus* genetic markers (*tlh* or *trh* or *tdh* or ratio of two markers, etc.) in the model. Water temp consisted of two measurements (surface water temperature and shore water temperature).

### Variable Generation

Three genetic variables consisting of the ratios *trh:tlh, tdh:tlh*, and *tdh:trh* were constructed to assess the relative abundance of the pathogenic markers comparatively as done previously in Flynn et al. (2019). All genetic variables, including pathogenic ratios, were log_10_ transformed. Time-lagged variables (e.g. air temperature measured 1-4 weeks prior to a given sampling event) were constructed and compared to time-indexed variables (i.e. at the time of sampling). For samples earlier in the season, lagged variables were treated as missing data. Correlation matrices were examined to determine redundancy in lagged and indexed variables. Earlier season entries for time-lagged variables were imputed through the multiple imputation methods used for missingness based on historic data from other seasons. In this way lagged entries were kept in the same sampling year and the imputed collection was in line with early/middle/late seasonal values.

Given the observed increase in total *V. parahaemolyticus* and pathogenic markers after prolonged exposure to air (Jones et al., 2016), an “exposure from surfacing” (EFS) variable was generated to characterize this association within the WDOH dataset. National Oceanic and Atmospheric Association (NOAA) tidal buoys and stations in Washington State were matched to WDOH sampling sites in order to determine the ambient air exposure times of sampled surfaced oysters. NOAA tidal data comprising of two pairs of high and low tide measurements per day were collected and aggregated using an application programming interface (NOAA, 2020). Tidal stations were matched to oyster sampling locations based on a non-Euclidean shortest water distance using the gdistance package in the R statistical computing environment (van Etten, 2017; R Core Team, 2020). The TideHarmonics package was used to interpolate tide height at all timepoints, assuming a mixed-Semidiurnal tidal cycle pattern (Stephenson, 2016).

All oyster samples were collected at 0.61 m (2 feet) above sea-level. EFS, which can also be described as the time a sample was exposed to ambient air before collection, was generated based on interpolated tidal heights. The variable was generated as follows:

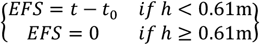

Where *h* is the tidal height at the time of sampling, t is the time of sampling, and t_0_ is the time that the oysters were exposed to ambient air.

### Model Development and Selection

The statistical analyses and modeling for this study were performed in R statistical and graphical software version 4.0.5 (R Core Team, 2021), modeling was performed with the *nlme* package (Pinheiro et al., 2021). Plots were created using the *ggplot2* package in R (Wickham, 2018), and maps were created using ArcGIS Pro software version 2.6.2.

Multiple imputation was used to impute data for: random missingness of environmental (9.66% missing) and genetic variables (19.04% missing), the nonrandom missing time-lagged data from early in the season, and missingness due to infrequent sampling at certain locations. To help improve model selection and performance. Absolute genetic variables were imputed using a censored imputation method first, which has been described previously (Davis et al., 2020). Using the *mice* package in the statistical software R (van Buuren and Oudshoorn, 2011), the overall dataset was imputed 100 times, each including ten iterations, using a multivariate imputation classification and regression tree (CART) model within years.

Mixed-effect time-series regression models were constructed to analyze the spatial and temporal relationship between environmental measures and abundance of *V. parahaemolyticus* genetic markers and their ratios (*tlh, trh, tdh, trh*:*tlh, tdh*:*tlh, trh*:*tlh*). Univariate and multivariate models were constructed based on trends observed in the exploratory data analyses. Separate sets of models were developed for each genetic marker and ratio that included environmental factors (including EFS) along with nested spatial and temporal dependence **(Figure 2)**. Univariate associations were first developed and followed by two- and three-way potential interactions between environmental factors, which were examined for inclusion or exclusion in multivariate models. Variance inflation factors were calculated to determine redundance in environmental variables.

Models including different environmental covariates, spatial and temporal random effects (intercepts and slopes), and time-lags were considered, including non-linear associations between the dependent and independent variables. Selection of final models was based on Bayesian Information Criterion (BIC) scores in addition to visual examination of covariate relationships. Variable imputation results were run in separate versions of each model. Model estimates, residuals, and BIC scores were pooled into a single model result, including fixed effect estimates and uncertainty and mixed effect variance (Grund et al., 2017). The *mitml* package in R was used to generate “pooled parameters” including imputation model summaries, estimates, and plots (Grund et al., 2019).

Residual temporal autocorrelation of residuals was detected in the models. To account for temporal autocorrelation, autoregressive– moving-average (ARMA) models were constructed for each of the six genetic outcomes based on the repeated weekly samples using the SSYG nested in zone model structure and were tested for lack of residual autocorrelation with a Breusch-Godfrey test (Breusch, 1978; Godfrey, 1978). In ARMA models, the AR (autoregressive) portion functions to estimate associations between the dependent variable and its own past values while the MA (moving average) element is comprised of modeling error as a grouping of contemporary errors and errors at various times in the past. Each model was fit with a modified Hyndman-Khandakar algorithm based on nested mixed-model structure (Hyndman and Khandakar, 2008), and each model’s residual temporal autocorrelation was examined through the autocorrelation function (ACF) and partial autocorrelation function (PACF). Models were examined pre- and post-application of ARMA correlation structure to determine existence of and remaining autocorrelation in the models. Data was checked for spatial autocorrelation via examination of semivariograms of data using SSYG’s as coordinates. The geographic distance matrix was constructed using the same non-Euclidian, water-distance algorithm to account for proximity based on watershed geography as in the construction of the *EFS* variable.

Several sensitivity regression analyses were conducted on early/late sample dates (i.e., those falling in May or October), sites with less than three samples in a year, and different ranges of years and specific oyster harvesting zones. Regressions using categorical variables of temperature quartiles were also run to look for nonlinear or dependent relationships between temperature and genetic variables. After controlling for all other effects, an existing wave pattern was discerned in the inter-year trend of *tdh*, which was fit to a sinusoidal function of the linear model as: *y* = *a*. sin(*x*) + *b*. cos(*x*).

## Results

The dataset of 2,836 samples were taken at 91 sampling sites located across 4 zones in Washington (**Figure 1**). There was a range of samples collected per year, with a minimum of 68 samples in 2005 and a maximum of 353 in 2007 (**Figure 3**). The majority (95.9%) of samples were collected between June-September, with the remaining samples collected in May or October (**Figure 4**). Sampling frequency by zone and year, with Hood Canal and Puget Sound being regularly sampled once and rarely twice per week, while Coastal Bays were sampled approximately every other week, and Northern Waters were sampled <1 per month (**Figure 4**). There were 288 unique SSYGs. Missing data (pre-imputation) was higher in the early and late weeks of the sampling season and was greater in the initial years of data collection (see **Figure S1** in the supplemental material). The Coastal Bays and Northern Waters zones had a higher percentage of missing values than Puget Sound or Hood Canal, and *tdh* was the most common missing data value.

**Fig. 3.**
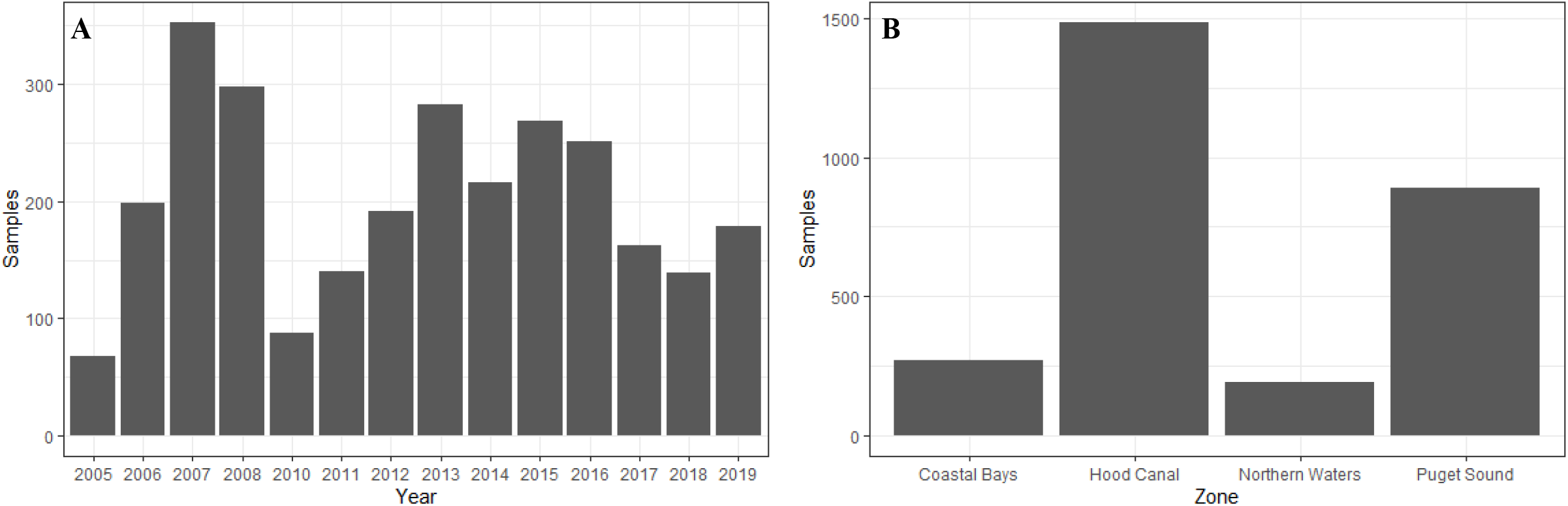
Cumulative oyster sampling reported by (A) year and (B).

**Fig. 4.**
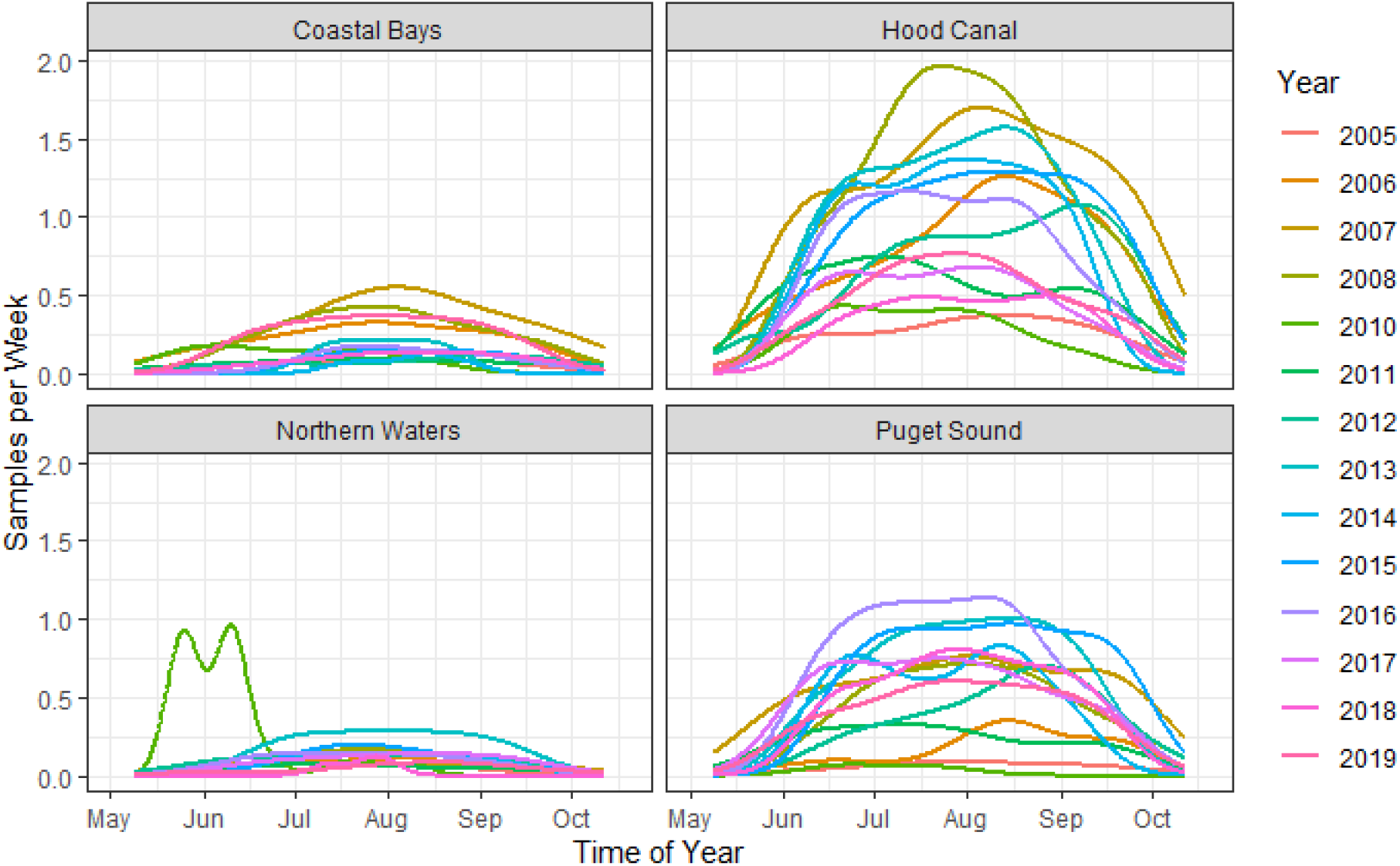
Kernel density plots of weekly sampling across time of year, stratified by zone and year. Note that the estimations are used solely as a visualization tool and not for statistical smoothing, as the number of samples and sampling frequency were not consistent across years.

The range of environmental variables were: ambient air temperature (5.2, 35.4 °C),surface water temperature (5.9, 30.6 °C), shore water temperature (9.9, 35.4 °C), tissue temperature (3.3, 38.4 °C), and salinity (0.6, 35.3 ppt) as seen in supplementary **Figures S2**. All genetic variables had values below the limit of detection (<0.3 MPN/g; (N_*tlh*_ = 9; N_*trh*_ = 101; N_*tdh*_= 469). Only *tlh* had values above the upper limit of quantification (N = 4) while *trh* had a maximum observable value of 46,000 MPN/g and *tdh* a value of 930 MPN/g. The *trh*:*tlh* and *tdh:tlh* ratios ranged from 0 to 100%, while *tdh*:*trh* ranged from 0 to 330%. Additional descriptions of the environmental and genetic variables are listed in **Table 1**.

**Table 1.** Descriptive characteristics of environmental and genetic variables including variation across sampling zones.

| Characteristic | Zones |  |  |  |  |
| --- | --- | --- | --- | --- | --- |
|  | Overall<br>(N= 2,836) | Coastal Bays<br>(N= 272) | Hood Canal<br>(N= 1,486) | Northern Waters<br>(N= 190) | Puget Sound<br>(N= 888) |
| Salinity (ppt) | 27.0 [5.88] | 29.0 [5.00] | 25.0 [8.00] | 27.2 [8.75] | 28.2 [4.00] |
| Tissue Temp (°C) | 20.0 [7.72] | 17.2 [4.50] | 20.4 [8.30] | 18.7 [4.71] | 21.1 [7.90] |
| Air Temp (°C) | 17.7 [5.90] | 16.1 [4.20] | 17.8 [6.00] | 17.0 [4.95] | 18.2 [5.83] |
| Shore Water Temp (°C) | 19.9 [4.70] | 19.4 [3.50] | 19.7 [4.80] | 21.6 [6.30] | 19.9 [4.60] |
| Surface Water Temp (°C) | 18.3 [3.50] | 17.8 [2.60] | 18.5 [4.10] | 17.3 [5.70] | 18.4 [3.00] |
| *tlh* (MPN/g) | 43.0 [426] | 4.2 [19.3] | 120 [921] | 4.3 [37.7] | 43.0 [421] |
| *trh* (MPN/g) | 4.2 [22.3] | 4.2 [14.1] | 3.80 [22.6] | 0.92 [3.84] | 4.30 [22.1] |
| *tdh* (MPN/g) | 0.3 [0.77] | 0.36 [0.77] | 0.3 [0.590] | 0.3 [0.59] | 0.36 [0.77] |
| *tdh:tlh* (ratio) | 0.01 [0.06] | 0.13 [0.36] | 0.004 [0.02] | 0.07 [0.10] | 0.01 [0.05] |
| *tdh:trh* (ratio) | 0.20 [0.50] | 0.130 [0.37] | 0.216 [0.97] | 0.363 [0.35] | 0.157 [0.38] |
| *trh:tlh* (ratio) | 0.057 [0.21] | 1.00 [0.59] | 0.025 [0.10] | 0.077 [0.10] | 0.084 [0.22] |

*Table displays median and inter-quartile range values for each characteristic of the overall dataset and stratified by harvesting zones.*

Air, tissue, and surface water temperatures showed similar seasonal (intra-annual) trends across years. Tissue temperature and air temperature were frequently higher in the Hood Canal and South Puget Sound. Surface water temperature displayed the same trend in all zones, although the Coastal Bays had cooler temperatures overall (**Figure S2**). Shore water temperature had a markedly different distribution, with no systematic variation in the Hood Canal and South Puget Sound between May-Aug. and then decreasing in Sep.-Oct; the Coastal Bays had an inverted relationship with temperatures dropping from May-Oct. Salinity was similar across years, outside of abnormally low salinity values in the first half of 2010. The Coastal Bays were the most saline overall (**Figure S2**).

Overall, *tdh* abundance was over an order of magnitude lower than *tlh* and was particularly low in Northern Waters; *trh* was similar but its abundance was higher than *tdh*. Unlike *tdh* and *trh, tlh* did not decrease in the later part of the growing season (Sep-Oct) (**Figure S3**). Negative trends for the genetic ratios were most prominent in the middle of the growing season, but the ratios were smaller in the early and late season, suggesting a reduced *tlh* abundance compared to *tdh* or *trh* in cooler temperatures (**Figures S4**).

All temperature measures were well-correlated with the highest correlations between tissue and air temp (r = 0.79), tissue and shore water temp (r = 0.76), and air and shore water temp (r = 0.68). All temperature measures were weakly positively correlated (r = 0.17-0.54) with *tlh, trh*, and *tdh*. Salinity was not correlated with temperature or any genetic variables. *tlh, trh*, and *tdh* were weakly positively correlated with one another. The genetic ratios were not correlated with temperature, salinity, or genetic variables (**Figures S5**). As previously noted in Flynn et al., 2019, we checked for shore and surface collinearity, which revealed a high VIF, and so the former was excluded from our models (**Figure S6**).

Exploratory analysis of ecological characteristics using LOESS identified non-linear relationships between *tlh* and *trh* with salinity and surface water temperature, and *tdh* with tissue temperature. Including these non-linear associations improved regression model fit for salinity (ΔBIC = 36.6), tissue temperature (ΔBIC = 20.5), and surface water temperature (ΔBIC = 40.7) compared to a model with only linear associations (**Figure S7**). Correlation matrices of lagged environmental characteristics did not display redundancy between time-indexed and lagged variables as shown in supplementary **Table S1**, and models’ BIC scores showed lagged salinity and air temperature measures to improve model fit when included in the model without their respective time-indexed values. For salinity in relation to the genetic markers, both 3- and 4-week lags improved model BIC scores but did not have a noticeable difference between them. Only the 3-week lag was chosen to have the simpler model. Similarly for air temperature, 1- and 2-week lags were identified by lower model BIC and 1-week lags were utilized for the same parsimonious modeling.

The exposure from surfacing variable EFS was not included in the final model as it was found to have no statistically significant association with any genetic marker in both univariate and multivariate models and did not demonstrate a mediating effect or interaction effect with any of the other environmental characteristics included in the model. Random slopes across independent variables were examined but were excluded due to model overfitting and failure of model convergence.

Fixed effects from regression models for *tlh* (a surrogate for total *V. parahaemolyticus* abundance), and pathogenic markers *trh* and *tdh*, are shown in **Table 2** and **Figure S8**. Each temperature variable displayed a statistically significant positive association in the univariate models for all three markers. Salinity at three weeks before the sampling event demonstrated a negative association with *tlh* and *trh* in the univariate models but not *tdh*. A non-linear association was observed between *tlh* and surface water temp where a statistically significant positive relationship began at 15°C and continued until 26°C before attenuating substantially. In contrast, there was a strong positive association between *trh* and surface water temperature, which was somewhat attenuated above 26°C. *tlh* and *trh* had non-linear relationships with salinity at 3-week lags, respectively, in which the negative relationship strengthened above 27 ppt. There was a threshold in the positive relationship between *tdh* and tissue temperature in that it was only observed above 20°C. All temperature associations became attenuated in the multivariate models for *tlh, trh*, and *tdh*. For surface water temp the trend observed was such that the association between both *tlh* and *trh* and surface water temperature above 26°C was no longer statistically significant. No statistically significant interaction was found between the temperature variables or with salinity (results not shown).

**Table 2.** Fixed effect estimates from univariate and multivariate regression models of *tlh, trh*, and *tdh* with environmental covariates.

|  | log *tlh* |  | log *trh* |  | log *tdh* |  |
| --- | --- | --- | --- | --- | --- | --- |
|  | Univariate | Multivariate | Univariate | Multivariate | Univariate | Multivariate |
| **Ecological characteristic** |  |  |  |  |  |  |
| Salinity (ppt) - 3 week lag | - | - | - | - | 0.00 (0.00, 0.01) | * |
| *1 ppt - 27 ppt* | -0.03 (-0.29, 0.23) | 0.04 (-0.17, 0.26) | -0.46 (-0.81, -0.11) | -0.37 (-0.67, -0.07) | - | - |
| *27 ppt - 35 ppt* | -0.61 (-0.91, -0.32) | -0.26 (-0.50, -0.02) | -0.90 (-1.32, -0.48) | -0.49 (-0.85, -0.13) | - | - |
| Tissue Temp (°C) | 0.07 (0.06, 0.08) | 0.02 (0.01, 0.03) | 0.09 (0.08, 0.1) | 0.04 (0.02, 0.05) | - | - |
| *3°C - 20°C* | - | - | - | - | 0.19 (-0.06, 0.45) | -0.03 (-0.31, 0.25) |
| *20°C - 38°C* | - | - | - | - | 1.24 (1.01, 1.48) | 0.75 (0.43, 1.06) |
| Air Temp (°C) - 1 week lag | 0.06 (0.05, 0.07) | 0.03 (0.02, 0.04) | 0.05 (0.04, 0.07) | 0.03 (0.02, 0.04) | - | - |
| Surface Water Temp (°C) | - | - | - | - | 0.06 (0.05, 0.08) | 0.02 (0.01, 0.04) |
| *6°C - 15°C* | -0.13 (-0.66, 0.40) | -0.17 (-0.64, 0.30) | 2.91 (2.51, 3.32) | 1.59 (1.15, 2.04) | - | - |
| *15°C - 26°C* | 2.21 (1.71, 2.71) | 1.40 (0.92, 1.89) |  |  | - | - |
| *26°C - 31°C* | 1.10 (0.11, 2.10) | 0.64 (-0.25, 1.52) | 2.06 (1.29, 2.83) | 1.01 (0.30,1.71) |  |  |

*Results are displayed as the log-transformed, pooled parameter estimates of the model with associated 95% confidence intervals. Reported associations are adjusted for region and year, as well as autocorrelation between weeks in the case of *tlh* and *tdh*. \* Indicated null effect (0) and exclusion from multivariate model.*

The ratios of *trh* in relation to total *V. parahaemolyticus* abundance (*trh*:*tlh*) and ratio of pathogenic strains in relation to each other (*tdh*:*trh*) had predictable associations with the environmental covariates given the associations observed for the absolute abundances. Therefore, the regression models for these two ratios are provide in **Table S2. Table 3** shows the fixed effects from the univariate and multivariate regression models for the ratio of *tdh* in relation to total *V. parahaemolyticus* abundance *(tdh:tlh*). In contrast to the previous multivariate models, associations with tissue temperature and air temperature were not statistically significant in any of the ratio models. Surface water temperature, however, continued to have a significant, non-linear relationship with *tdh:tlh* but not with *tdh:trh* or *trh:tlh*.

**Table 3.**
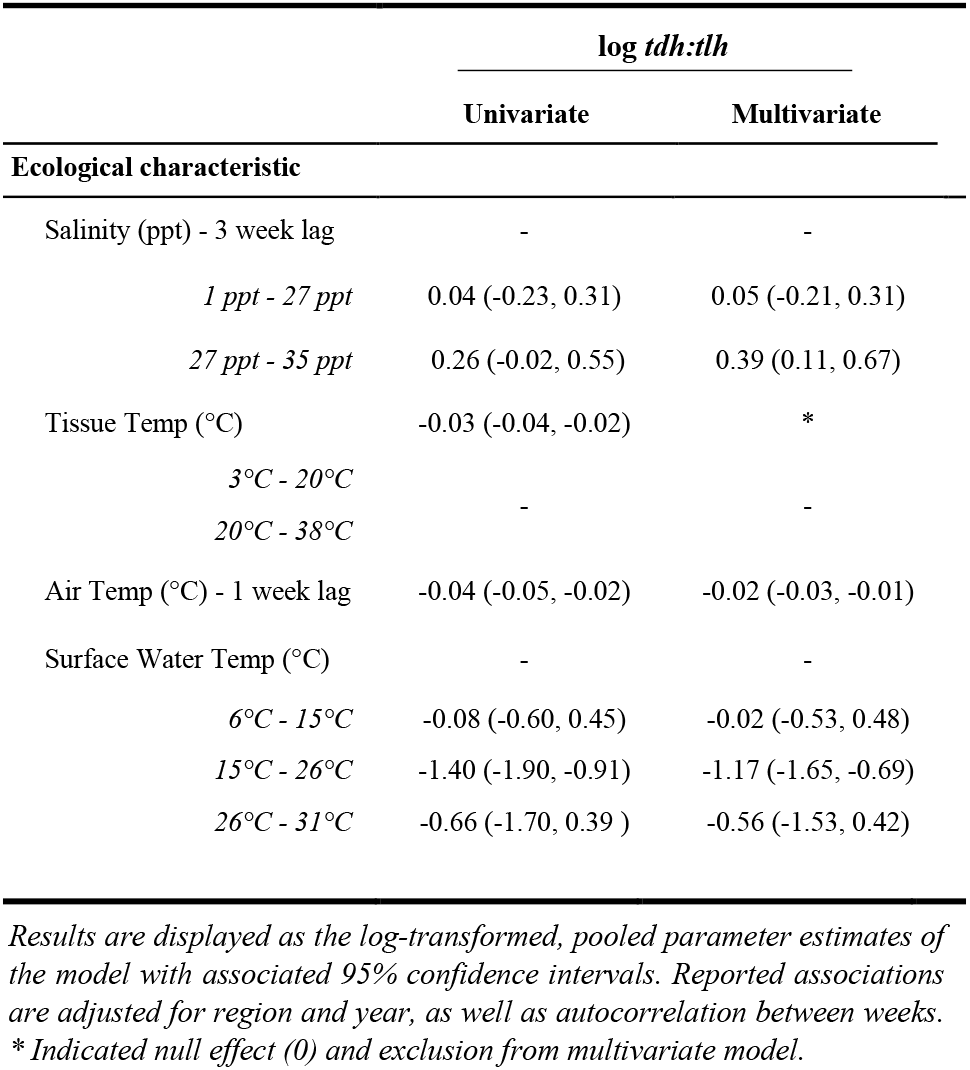
Univariate and multivariate associations between the *tdh:tlh* ratio with environmental covariates.

The random effects for all univariate and multivariate models are reported in **Table 4** and supplementary material **Table S3**. The random effect for year was redudant with the inclusion of the SSYG random effect, and a nested region/SSYG random effect structure had the best model performance with the lowest BIC scores. There was minimal variation across SSYGs compared to zones, except for *tdh* (and to a lesser extent *trh*) where the random intercepts for both SSYG and zone were similar. When a temporal random intercept for week was incorporated into the models it resulted in overfitting, as samples were taken roughly a week apart and the models were singular. While random slopes for the environmental covariates were considered, none resulted in any noteworthy changes to model interpretation.

**Table 4.** Model Random Effects for Zone and Sample Site Year Group (SSYG)

|  | log *tlh* | log *trh* | log *tdh* | log *tdh:tlh* |
| --- | --- | --- | --- | --- |
| Random Effects |  |  |  |  |
| Univariate |  |  |  |  |
| Salinity (ppt) |  |  |  |  |
| *Random Intercept - Zone* | 0.33 | 0.05 | 0.04 | 0.26 |
| *Random Intercept - SSYG* | 0.41 | 0.13 | 0.14 | 0.44 |
| Tissue Temp (°C) |  |  |  |  |
| *Random Intercept - Zone* | 0.25 | 0.02 | 0.03 | 0.24 |
| *Random Intercept - SSYG* | 0.39 | 0.17 | 0.14 | 0.44 |
| Air Temp (°C) |  |  |  |  |
| *Random Intercept - Zone* | 0.31 | 0.04 | 0.03 | 0.26 |
| *Random Intercept - SSYG* | 0.40 | 0.14 | 0.14 | 0.45 |
| Surface Water Temp (°C) |  |  |  |  |
| *Random Intercept - Zone* | 0.32 | 0.12 | 0.04 | 0.26 |
| *Random Intercept - SSYG* | 0.33 | 0.17 | 0.16 | 0.38 |
| Multivariate |  |  |  |  |
| *Random Intercept - Zone* | 0.22 | 0.06 | 0.03 | 0.19 |
| *Random Intercept - SSYG* | 0.00 | 0.01 | 0.05 | 0.02 |

Estimate of random intercept effect size for univariate and multivariate models. Sample Site Year Groups were time and space matched locations such as Totten Inlet-2017.

Residuals of each of the models showed statistically significant autocorrelation across weeks nested within years in both the ACF/PACF plots and by the Breusch-Godfrey test; therefore, each model displayed in Tables 2 & 3 was further fit in a sensitivity analysis with autoregressive and moving-average terms (ARMA) (**Table 5** and **Figure S9**). Applying residual structure reduced model residual autocorrelation for each model but did not show change in regression associations. Residual semivariograms did not indicate any spatial dependence **(Figure S10)**. Given that exploratory analyses identified *tdh* abundance oscillating across years, another sensitivity analysis fit a sinusoidal trend to the yearly variation of *tdh* adjusting for previously included environmental characteristics (**Table 6)**. Model fit improved with the addition of the sinusoidal parameters to the model (ΔBIC = 16), with a wave period of approximately 15 years (**Figure 5)**.

**Table 5.** Model variable temporal autocorrelation structure and residual significance

| Variable | ARMA (p,q) model |  | Breusch–Godfrey test |  |
| --- | --- | --- | --- | --- |
|  | Coefficient | Std. Error | BG | P-value |
| *tlh* (MPN/g) | - | - | 4.58 | 0.21 |
| AR(1) | 0.79 | 0.02 | - | - |
| MA(1) | -0.38 | 0.03 | - | - |
| *trh* (MPN/g) | - | - | 3.20 | 0.36 |
| AR(1) | 0.74 | 0.04 | - | - |
| MA(1) | -0.40 | 0.05 | - | - |
| *tdh* (MPN/g) | - | - | 2.72 | 0.44 |
| AR(1) | 0.79 | 0.03 | - | - |
| MA(1) | -0.56 | 0.04 | - | - |
| *trh:tlh* | - | - | 0.56 | 0.91 |
| AR(1) | 0.92 | 0.02 | - | - |
| MA(1) | -0.65 | 0.03 | - | - |
| *tdh:tlh* | - | - | 1.72 | 0.63 |
| AR(1) | 0.85 | 0.02 | - | - |
| MA(1) | -0.52 | 0.03 | - | - |
| *tdh:trh* | - | - | 4.81 | 0.19 |
| AR(1) | 0.85 | 0.03 | - | - |
| MA(1) | -0.60 | 0.05 | - | - |

Reported associations are adjusted for all environmental and genetic covariates as well as region and year. Nested ARMA model of multivariate models’ residuals for one (1) time step. Nonsignificant p-value indicates lack of statistical significance of autocorrelation of linear model error term.

**Table 6.** Univariate and multivariate sinusoidal function of *tdh* across years.

| Model Coefficients | Univariate | Multivariate |
| --- | --- | --- |
| $\sin(2 * \pi * (t / \text{total } t \text{ in years}))$ | 0.19 (0.11, 0.27) | 0.20 (0.12, 0.28) |
| $\cos(2 * \pi * (t / \text{total } t \text{ in years}))$ | 0.14 (0.06, 0.23) | 0.06 (-0.03, 0.15) |

**Fig. 5.**
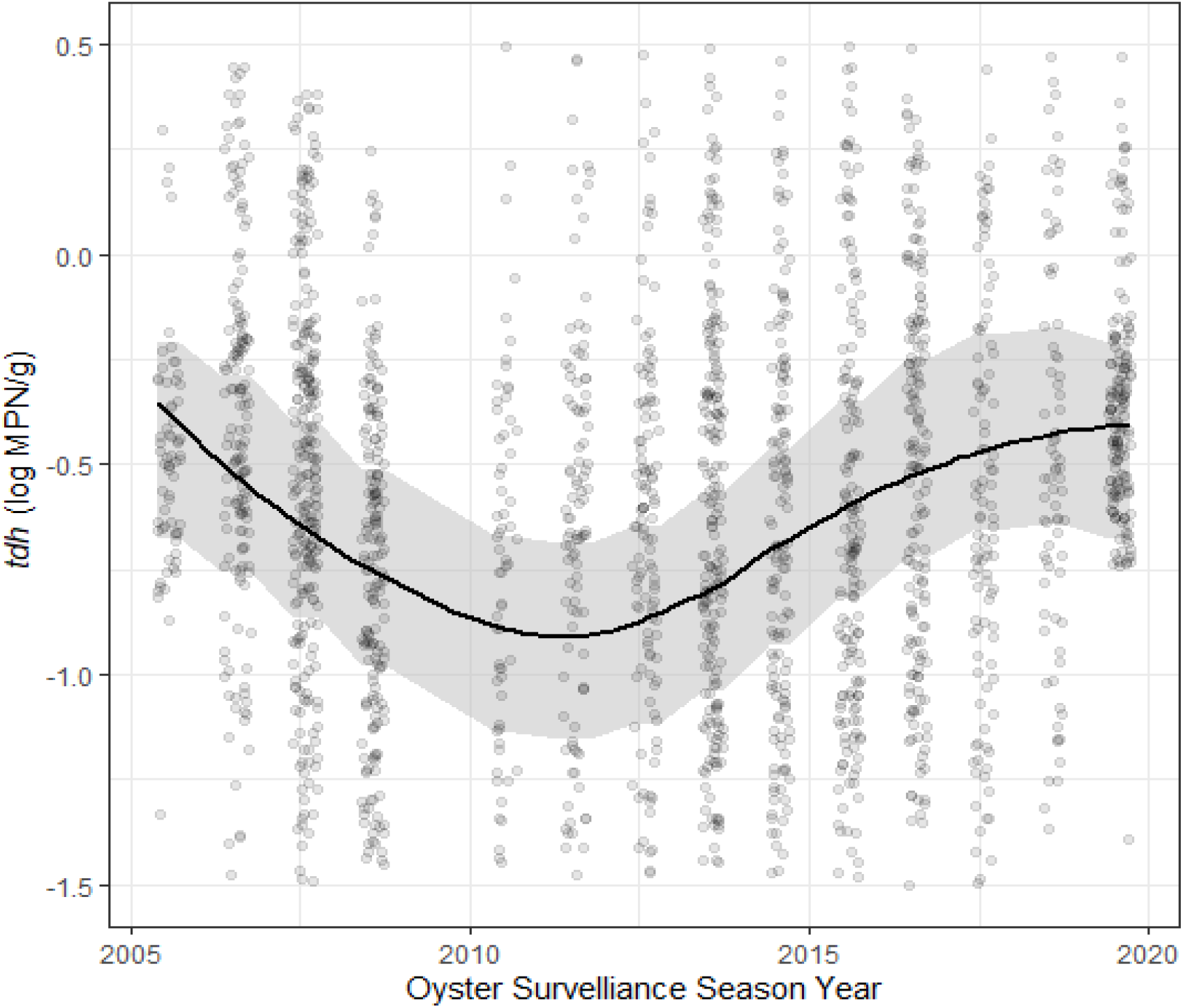
Sinusoidal trend of log *tdh* values across time. Plotted results of univariate model coefficients shown in Table 6.

Sensitivity analyses removing and including specific years of sampling revealed that for models using only post-2013 data there was a more pronounced negative association between salinity and *tlh* and the positive association between tissue temperature and *tdh* strengthened. Early and late season exclusions did not substantially change model associations. Restricting the dataset to only repeatedly sampled SSYG’s resulted in a significant attenuation in the negative relationship between salinity and *trh* but a slight strengthening in the negative slope of the relationship between salinity and *tlh*. Restricting model associations by harvesting zone revealed difference in the effect size of salinity on the model, with a noticeable difference between the Coastal Bays and Northern Waters zones, and the Hood Canal and Puget Sound zones.

## Discussion

This analysis evaluated the relationships between environmental measures and genetic markers of *V. parahaemolyticus* using a large number of samples of Pacific oysters in the U.S. state of Washington. The analysis accounted for hierarchical spatial and temporal heterogeneity by including random effects in regression models. Several non-linear relationships were observed between environmental characteristics and *V. parahaemolyticus* genetic markers. The nested models created from the extensive WDOH dataset complemented by multi-step imputation indicate that there are differences in the magnitude of the associations for *tlh* and *tdh*:*tlh* across harvest zones and that genetic marker abundance experiences temporal autocorrelation (i.e., abundance from previous weeks is indicative of abundance at the time of sampling). The nature of the relationships did not differ by zone. The utility of time-lagged measures was demonstrated in the modeling of the relationship between salinity and air temp and *tlh* and *trh*, but not *tdh*. Overall, these models provide greater insights into the variation of each of the genetic markers for *V. parahaemolyticus* in Washington, the potential larger ecological conditions that may be affecting them, and the complexity of the estuarine ecosystems that *V. parahaemolyticus* and the pacific oyster inhabit in Washington state.

The constructed exposure from surfacing (EFS) variable was not associated with the abundance of any genetic marker. This was unexpected, since exposure to relatively warmer air and direct sunlight have been show in experimental settings to increase the abundance of *Vibrio* spp. in oysters (Nordstrom et al., 2004; Jones et al., 2016). This null association is almost certainly not an indication of exposure time not increasing *V. parahaemolyticus* growth but instead likely due to NOAA tidal data not capturing the specific tidal fluctuations of the numerous inlets in the Washington state oyster growing environment. The wide flat beaches have significant variation in where and how long oysters are exposed, and sample oyster gathering is not precise enough at this time to ensure oysters were sampled exactly when completely exposed, or at the same point on the same site every time. Incorporating specific, recurring sampling coordinates for sample sites and the use of reliable tidal dataloggers to monitor submersion and exposure cycles more precisely would likely better capture the true relationship of exposure time and *V. parahaemolyticus* abundance accurately. The EFS variable did not account for weather patterns as it was prohibitively difficult to retrieve and summarize cloud cover data for the entire study area across all sampled summers. However, the modifying effect of solar radiation on oyster surfacing time likely plays an important role in *V. parahaemolyticus* growth in oyster tissue (USDA, 2005). Accounting for weather patterns in future analyses may alter the null association observed with EFS.

Tissue temperature exhibited a strong positive, albeit non-linear association with *tdh*, with no significant association observed below 20°C. As post-harvest oyster tissue temperature controls are a key piece of WDOH’s vibriosis control plan, identification of such a threshold for pathogenic growth should be examined further (Nilsson et al., 2019). An upper threshold limit for the association between surface water temperature and *tlh* was also reported in an analysis of a subset of the dataset used in this study (Flynn et al., 2019); however, the full dataset identifies the threshold at a higher temperature (26°C vs. 22°C). This difference could be due to the imputation of previously missing temperature values in this analysis, as the range of temperature values was extended in the complete dataset compared to previous subset of data that excluded sampling occasions with any missing data. A previously unidentified lower threshold in the *tlh*-surface temperature association was also observed around 15°C, which is consistent with the minimum required temperature for *V. parahaemolyticus* growth reported in laboratory and field settings (Urquhart et al., 2016; Wang et al., 2018). The positive association between surface water and *trh* also demonstrated an upper threshold at 26°C but with no lower threshold. This may be because *V. parahaemolyticus* strains that contain the *trh* marker continue to decline as water temperatures drop. The larger association between *trh* and water temp compared to *tlh* indicates a risk of pathogenic *V. parahaemolyticus* having more rapid growth rates in warmer waters when compared to non-pathogenic bacteria. Notably, the relationship between surface water temperature and each of the genetic markers did not change in sensitivity analyses stratified by the four growing zones. This suggests consistency in this relationship across these ecologically diverse waters which was possibly due to random intercepts accounting for baseline differences in genetic targets. The inconsistency between strains of *V. parahaemolyticus* containing the *tdh* marker compared to *tlh* or *trh* in relation to water temperature had not previously been observed (Flynn et al., 2019).

Salinity (lagged at 3-weeks) also demonstrated a non-linear negative relationship with *tlh* and *trh* with a threshold at 27 ppt, where the magnitude of the relationship strengthened and in the case of *tlh* became statistically significant. This non-linear relationship supports previous findings of a negative relationship between salinity and *V. parahaemolyticus* at high (greater than 23 ppt) salinity levels (Davis et al., 2017). Although previous studies in Washington did not observe an association with salinity (Flynn et al., 2019; Paranjpye et al., 2015), complex non-linear relationships with salinity and *V. parahaemolyticus* have been found in other bodies of water (DePaola et al., 2003; USFDA, 2005; Johnson, 2015; Davis et al., 2017; Martinez-Urtaza et al., 2016). Further, the previous PNW studies only considered salinity measurements at the time of sampling, whereas statistically significant relationships in this study were identified using a time-lagged measure of salinity. This suggests that the salinity of the harvesting water weeks before sampling may impact growth of *V. parahaemolyticus* in microbial ecological communities, possibly due to seawater intrusion diluting the existing population. The significant negative effect on overall abundance in saline waters over 27 ppt is in line with similar previous estimates (DePaola, et. al., 2000; Martinez-Urtaza et al., 2016) but our results show this relationship’s threshold varies by the zonal environment of the oyster.

The temporal autocorrelations of each genetic marker and ratio were estimated using an ARMA model to account for any stationary stochastic process. The significant results at AR(1) and MA(1), each to the order of one time-step, indicate that the concentration of genetic markers at a specific site in a particular year are influenced by some unobserved moving average (likely a seasonal trend) as well as previous concentrations from that site (the autoregression). This outcome suggests a temporal relationship process of previous week’s influence on *V. parahaemolyticus* abundance for the current week.

The sinusoidal curve function of *tdh* across years implies that some other long-term ecological process is not being captured by the measured environmental variables included in this analysis. The full sampling period of ∼15 years would potentially indicate this trend was tied to a longer-time scale process such as the El Niño–Southern Oscillation (ENSO) (Logar-Henderson et al., 2019) which it superficially resembles. Previous studies have linked the dissemination of pathogenic *V. parahaemolyticus* to the expansion and dynamics of the poleward propagation and receding of El Niño ocean water in South America (Martinez-Urtaza et al., 2008). This relationship could be due to increased precipitation brought in ENSO years, but more research is needed to identify what process is taking place. Anomalously high heat led to a vibriosis outbreak in the PNW during the summer of 2020. Unfortunately, sampling in Washington was heavily impacted by the pandemic and so was not included in the current dataset. The models developed in this analysis can be used to extrapolate to high temperatures in an attempt to predict *V. parahaemolyticus* levels in future climatic scenarios. The sinusoidal curve function observed for *tdh* abundance also appears to have predicted a high level of *V. parahaemolyticus* in 2020 and so may also inform future large-scale trends.

A limitation of this study was that although generally sampling at sites occurred no more than once a week, sampling intervals usually ranged between 3-12 days, which may have introduced some amount of measurement error. Therefore, differences in the “optimal” week lags should be understood to not specifically refer to 7-day increments. Oyster to oyster variability could also alter abundance estimates (e.g., if a single oyster remained inactive over a period of exponential growth). However, such variability should be random and would therefore only bias the observed associations towards the null. The WDOH sampling design limited our ability to effectively isolate the temporal random effect size for individual years in our models. Each year, WDOH would move sampling locations, sometimes clustering in a specific region in successive years and sometimes moving significant distances. To account for this spatial “drift” we constructed the SSYG variable to assign sampling locations to consistent sites within a year. When incorporating nested random effects however, the SSYG confounded the year random effect and prevented us from including both sets of random effects in the models. However, as SSYG accounted for the year random effect, this created a more parsimonious model.

The associations described in this work are notable, but further environmental information, including measurements of water quality (e.g., oxygenation, turbidity, plankton abundance, suspended solids), watershed precipitation, and land use, were not readily obtainable to include in the analysis. This missing information could explain some of the associations observed in this work, possibly the residual temporal variation in the models (Paranjpye et al., 2015; Davis et al., 2017; Nilsson et al., 2019). This information has been used for analyses of *Vibrio* spp. in the Chesapeake Bay and other regions and future work could incorporate such measurements if they could be regularly gathered during oyster sampling (DeLuca et al., 2019; Davis et al., 2017; Johnson et al., 2012). Complementary datasets of these variables collected at different times and sampling locations could also be incorporated into a spatiotemporal prediction framework in order to provide additional inputs into the models described in the current work. This framework can also be used to make more informed imputations for missing data in the existing WDOH shellfish dataset. These potential models could further utilize spatial–temporal statistics to forecast *V. parahaemolyticus* abundance in Washington State, providing a resource shellfish harvesters and risk managers can use to make informed food safety decisions.

Modernizing the FDA’s *V. parahaemolyticus* risk assessment based on improved statistical and analytical tools with more granular measurements of *V. parahaemolyticus* samples and environmental conditions, such as was done in this study, would allow for increasingly proactive risk management options. Regional heat anomalies, like that observed in the PNW in the summer of 2020, often drive risk and are linked to the most severe vibrioses outbreaks. The FDA *V. parahaemolyticus* risk assessment was intended to predict regular, sporadic cases and an important extension of the assessment would be to forecast outbreaks (USFDA, 2005). The findings in the current study and other recent studies should be compared to the *V. parahaemolyticus* risk assessment assumptions and estimates in order to create a more perfect tool to manage pathogenic *V. parahaemolyticus* risk. Application of spatial imagery or dynamic and interactive models such as those utilized in the latest norovirus risk assessment (Pouillot et. Al, 2021) could be helpful for adapting the modeling approach of this paper to a future interactive *V. parahaemolyticus* risk assessment tool. Overall, increased focus on updating shellfish safety assessments in the PNW using new methods and advanced models is of increasing importance given more frequent intense heat events along with the rise in shellfish borne illnesses.

## Conclusion

Modeling the abundance of genetic markers of total *V. parahaemolyticus* (*tlh*) and potentially pathogenic strains (*trh, tdh*) in Pacific oysters from Washington State revealed strong associations with surface water temperature and salinity. Overall, this study confirmed the existence of a positive trend between water temperature and *tlh* with an upper threshold while also identifying a previously unobserved lower threshold. *trh* appeared to have a similar relationship with salinity and temperature to *tlh* while *tdh* had a positive relationship with tissue temperature in warmer oysters (>20°C). This study also identified an interannual sigmoidal curve for *tdh*, suggesting long-term ecological variation that may impact the risk of vibriosis to oyster consumers. These findings show that mixed models incorporating spatial and temporal variation can reveal the intricate links between environmental measures and the potential growth of pathogenic strains of *V. parahaemolyticus*. We recommend that subsequent models to explain *V. parahaemolyticus* estimations incorporate geostatistical techniques in order to identify zonal and sub-zonal differences across shellfish growing environments to better estimate the risk of shellfish-borne illness among consumers of Pacific oysters

## Acknowledgements

The authors would like to thank Laurie Stewart (Office of Communicable Disease Epidemiology), Gina Olson (Public Health Laboratory), and Lawrence Sullivan (Office of Environmental Health and Safety) from the Washington State Department of Health for their collaboration and assistance in creating, organizing, and maintaining the data used in this analysis. Additional thanks are given to the countless state and local staff and student interns who were involved shellfish sampling, laboratory analysis, and illness investigations. Finally, thanks are given to Tim Shields from the Spatial Science for Public Health Center for helping to create the maps displayed in this paper.

## Data Availability Statement

The datasets generated for this study are available on request to the corresponding author.

