## Supplemental Figures and Tables for "Nested spatial and temporal modeling of environmental conditions associated with genetic markers of *Vibrio parahaemolyticus* in Washington state Pacific oysters"

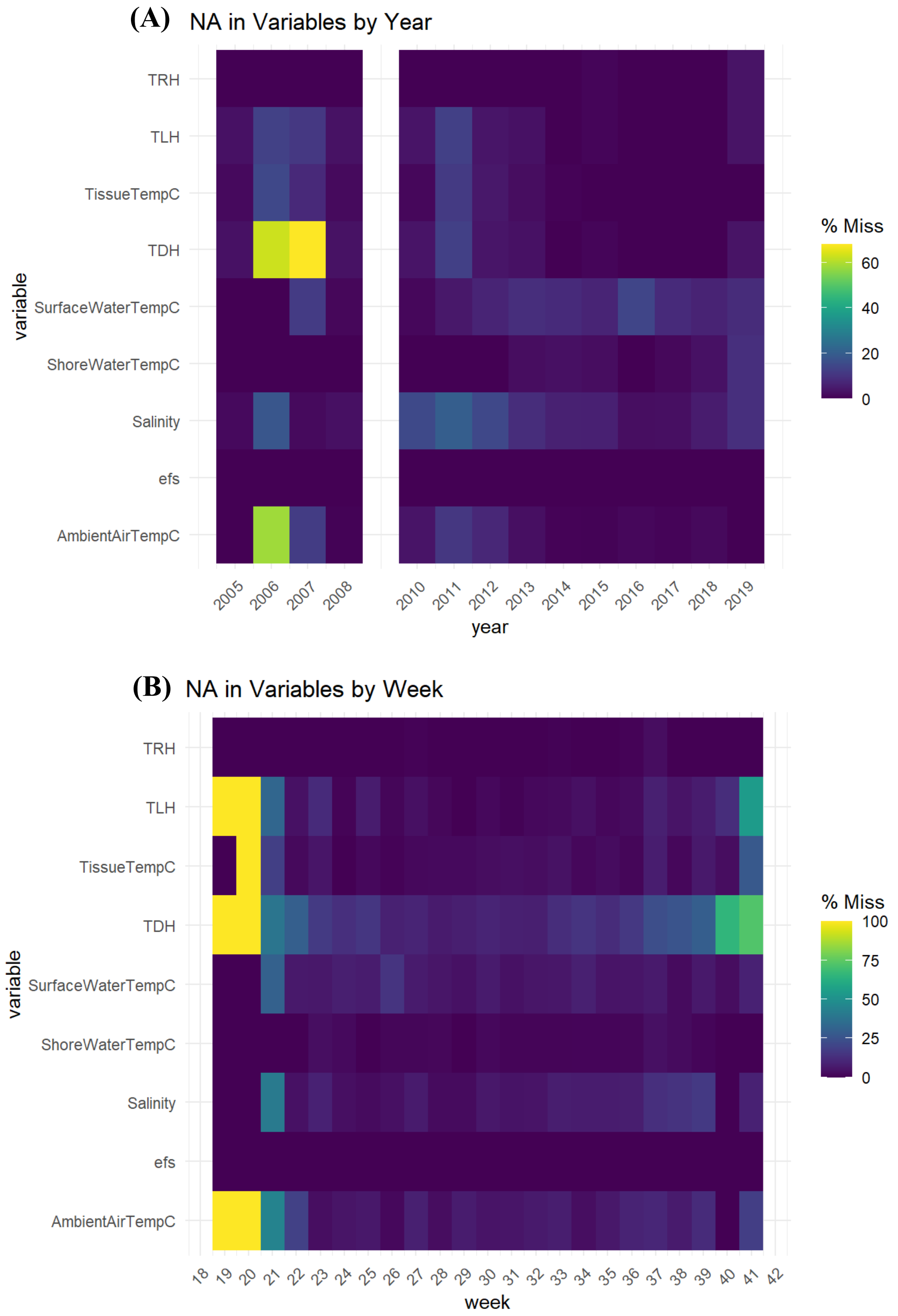

**(C)**

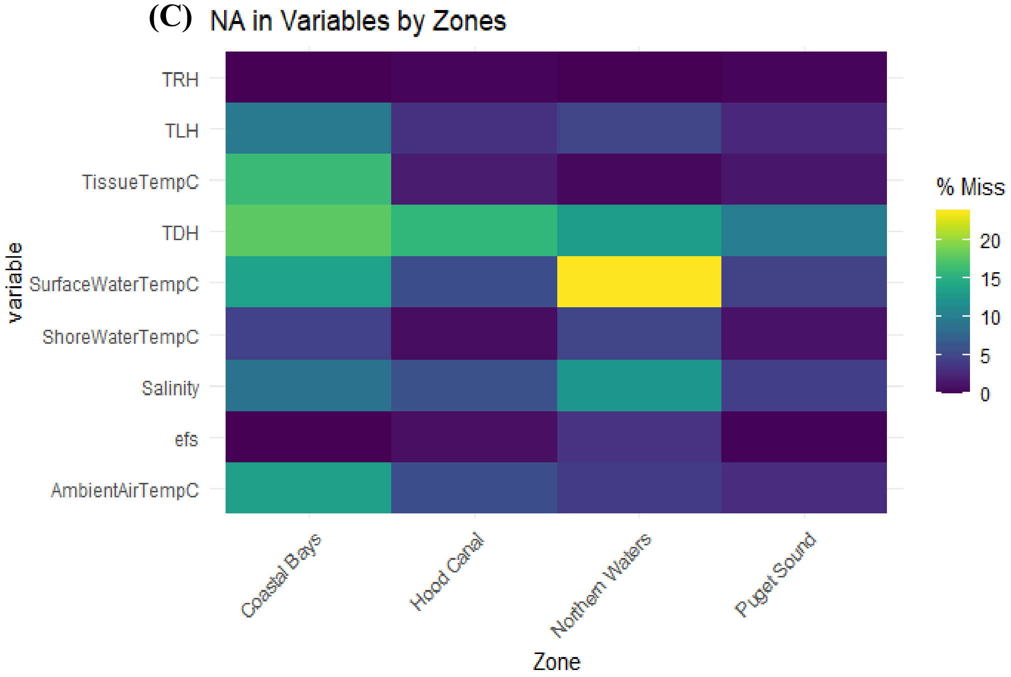


**Figure S1.** (A): Pre-Imputation Dataset Missingness by year (2009 excluded). (B): Pre-Imputation Dataset Missingness by sampling week in year. (C): Pre-Imputation Dataset Missingness by sampling Zone.

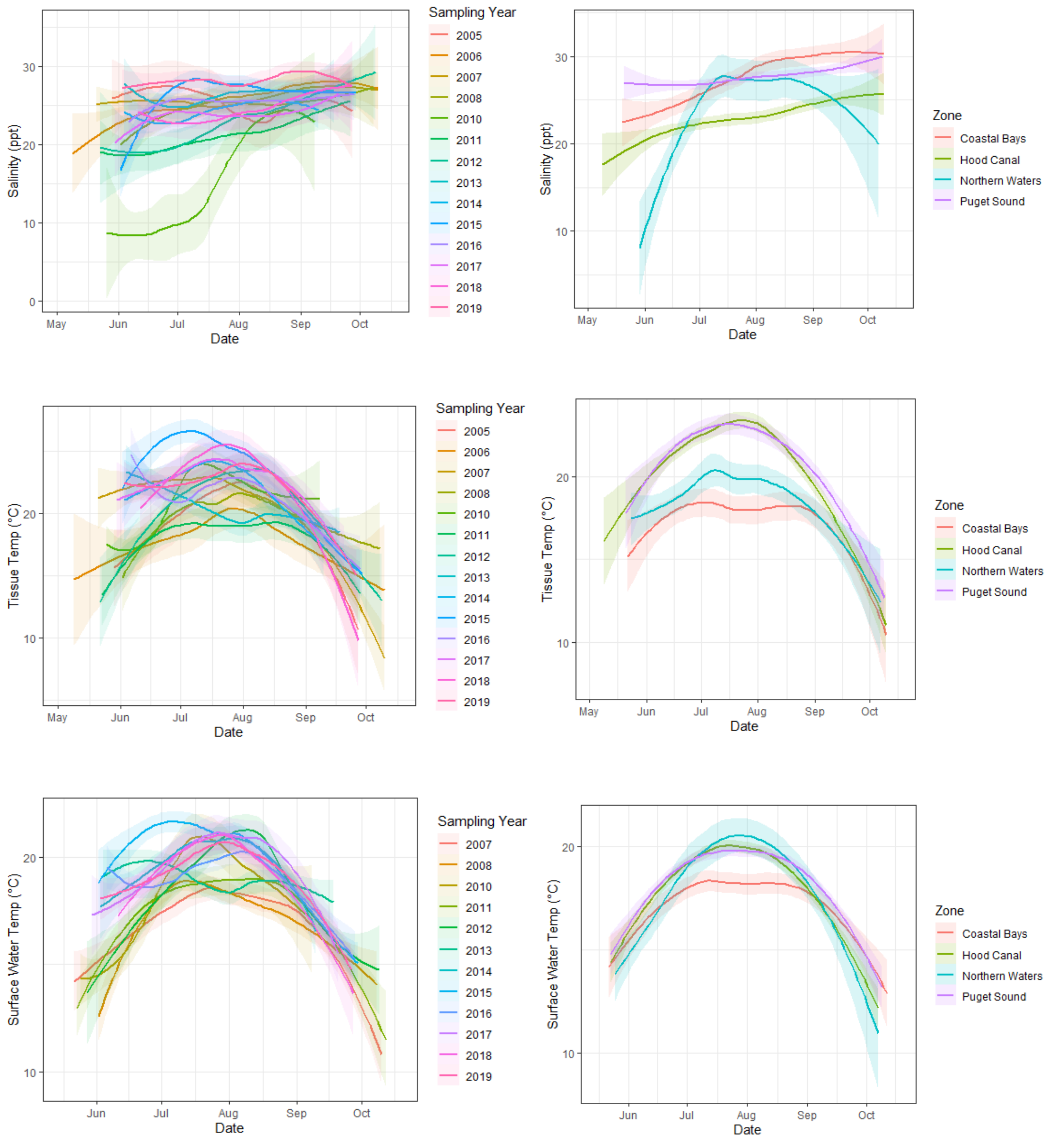


**Figure S2**. LOESS plots of environmental characteristics over intra-annual time, stratified by year and zone.

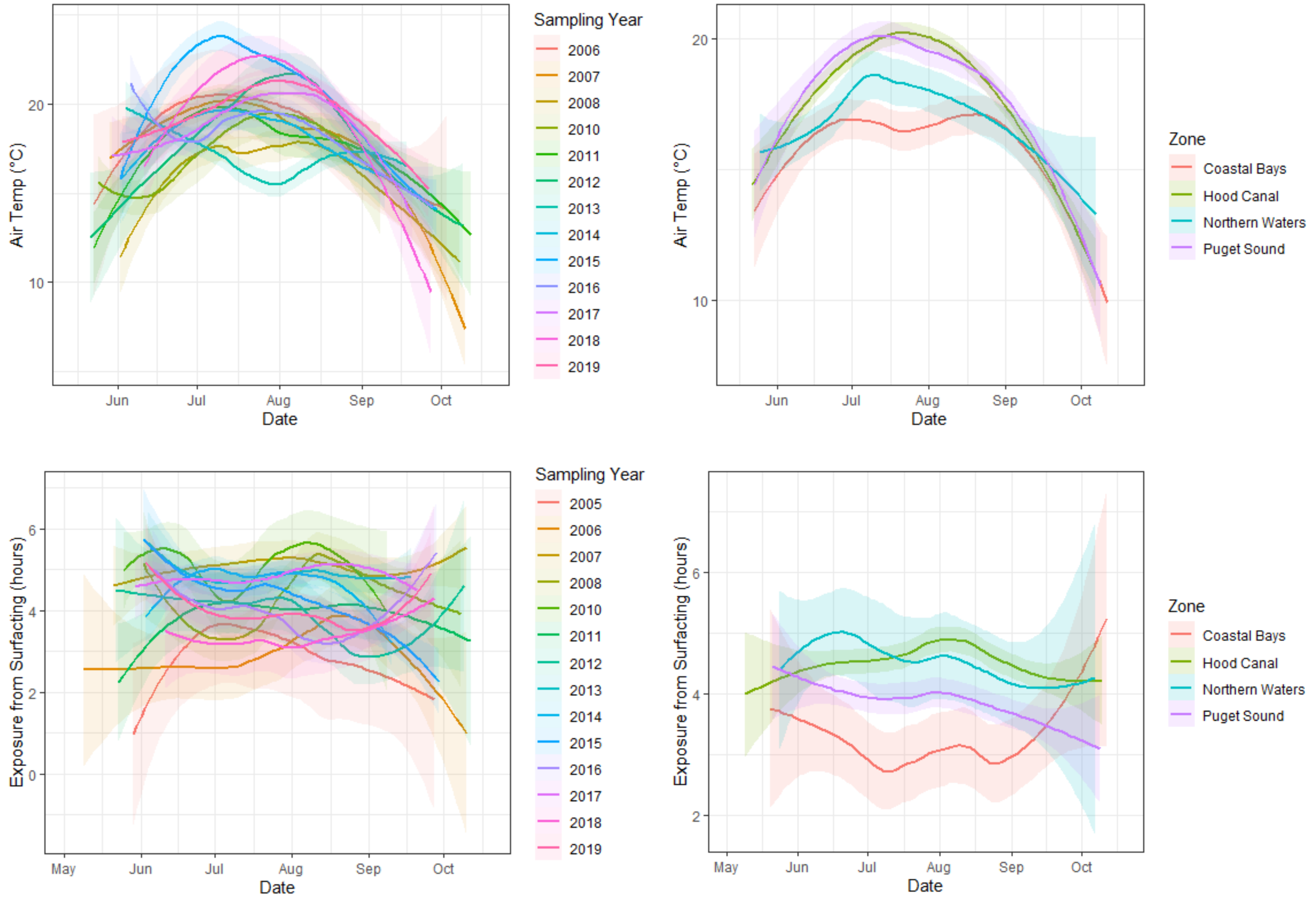


**Figure S2 cont**. LOESS plots of environmental characteristics over intra-annual time, stratified by year and zone.

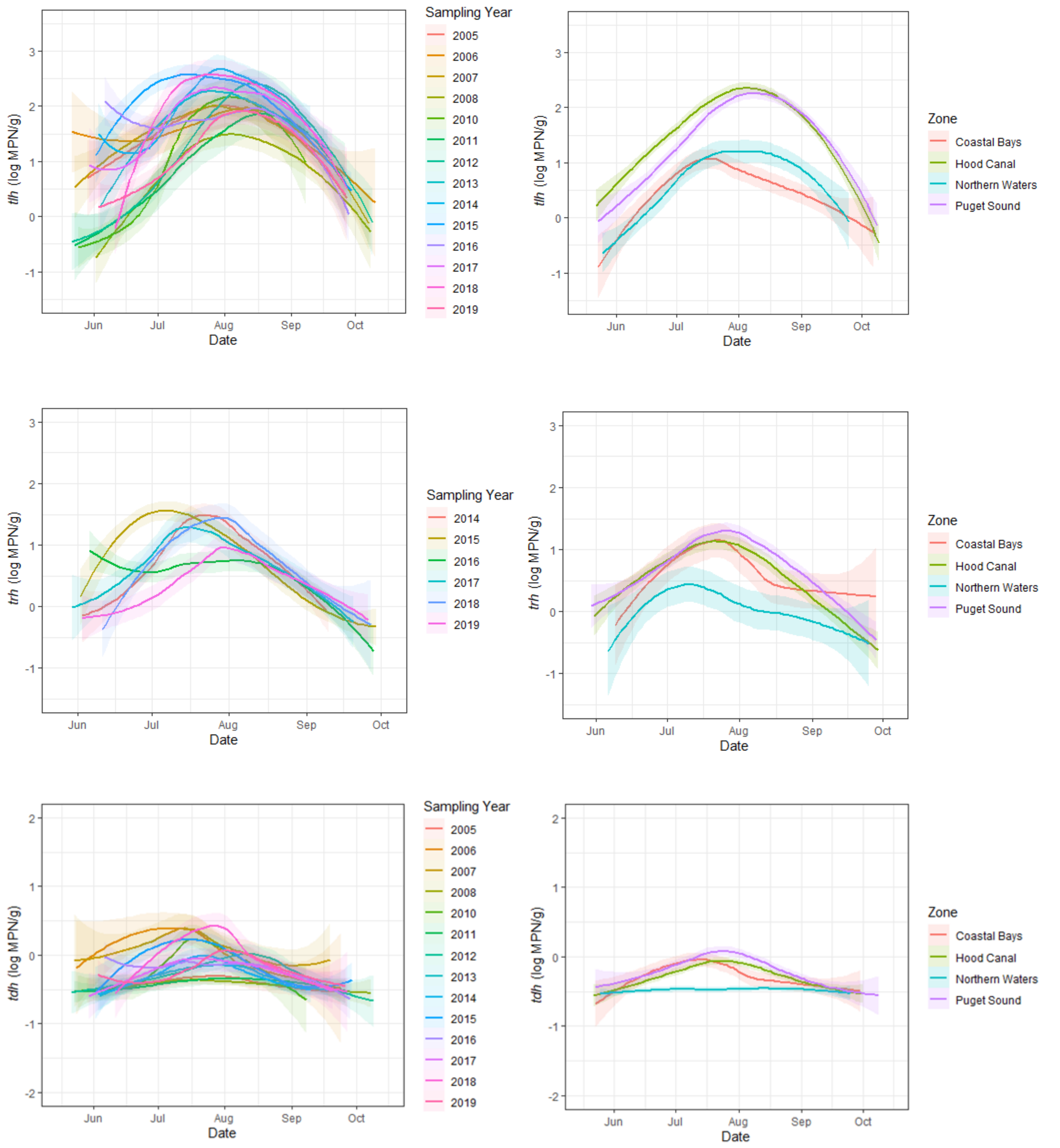


**Figure S3**. LOESS plots of *V. parahaemolyticus* genetic markers over intra-annual time, stratified by year and zone. Plots display a representative imputation for samples below and above the limit of detection.

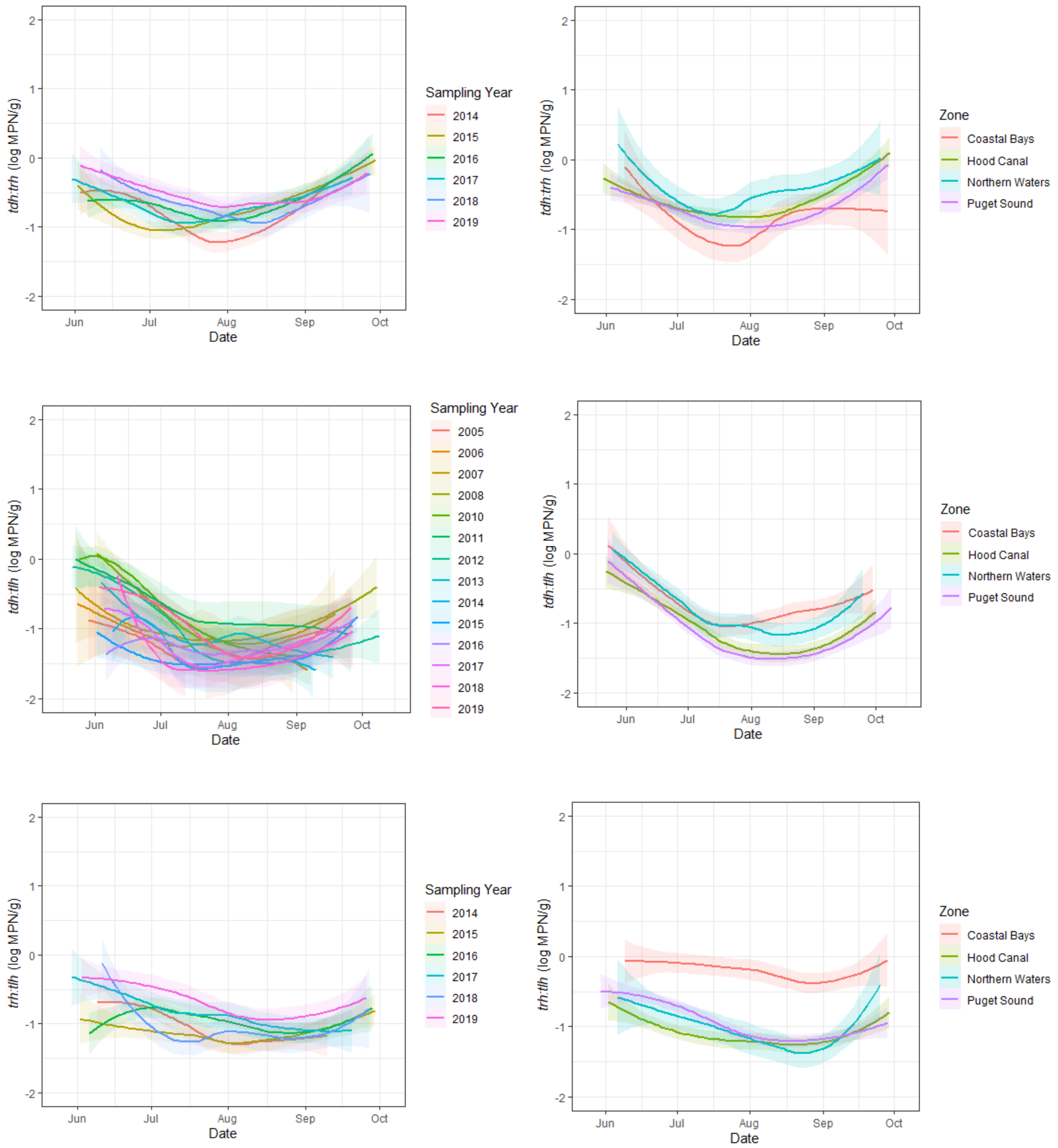


**Figure S4**. LOESS plots of *V. parahaemolyticus* genetic marker ratios over intra-annual time, stratified by year and zone. Plots display a representative imputation for samples below and above the limit of detection.

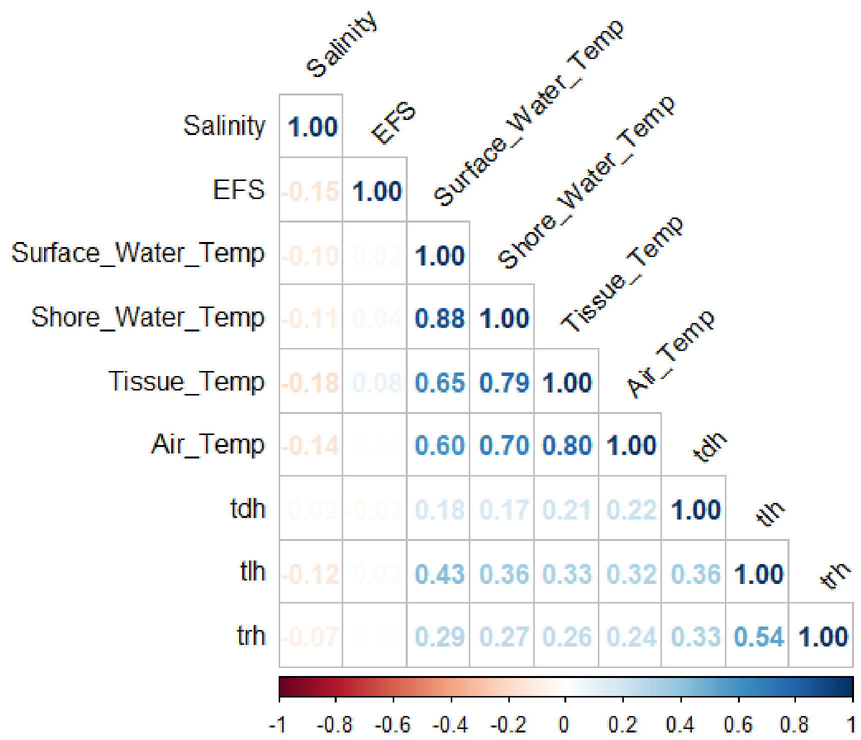


**Figure S5.** Correlation matrix of covariate values

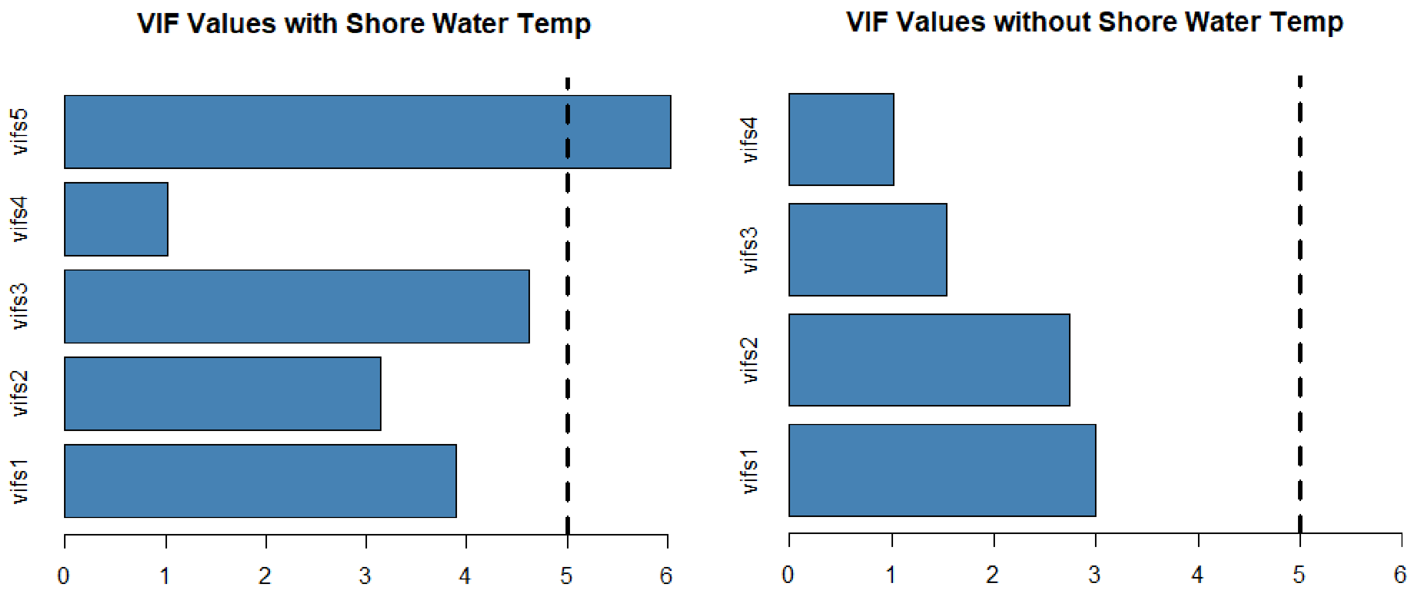


**Figure S6.** Variance inflation factors of model including and excluding shore water temperature. Variables vifs1 = tissue temperature, vifs2 = Air temperature, vifs3 = surface water temperature, vifs4 = salinity, vifs5 = shore water temperature.

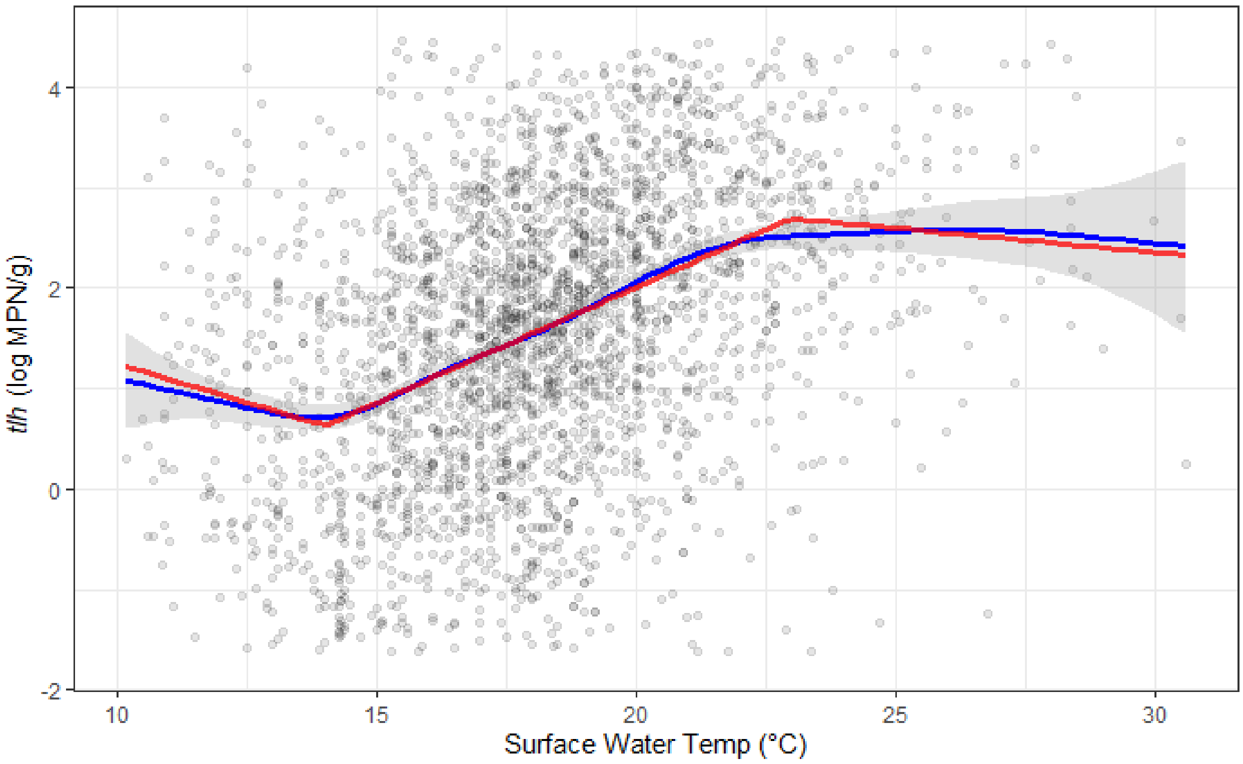


**Figure S7A.** Comparison of univariate regressions for surface water temperature to *tlh* with linear regression with spline term (red) compared to smoothed conditional mean line of data (blue).

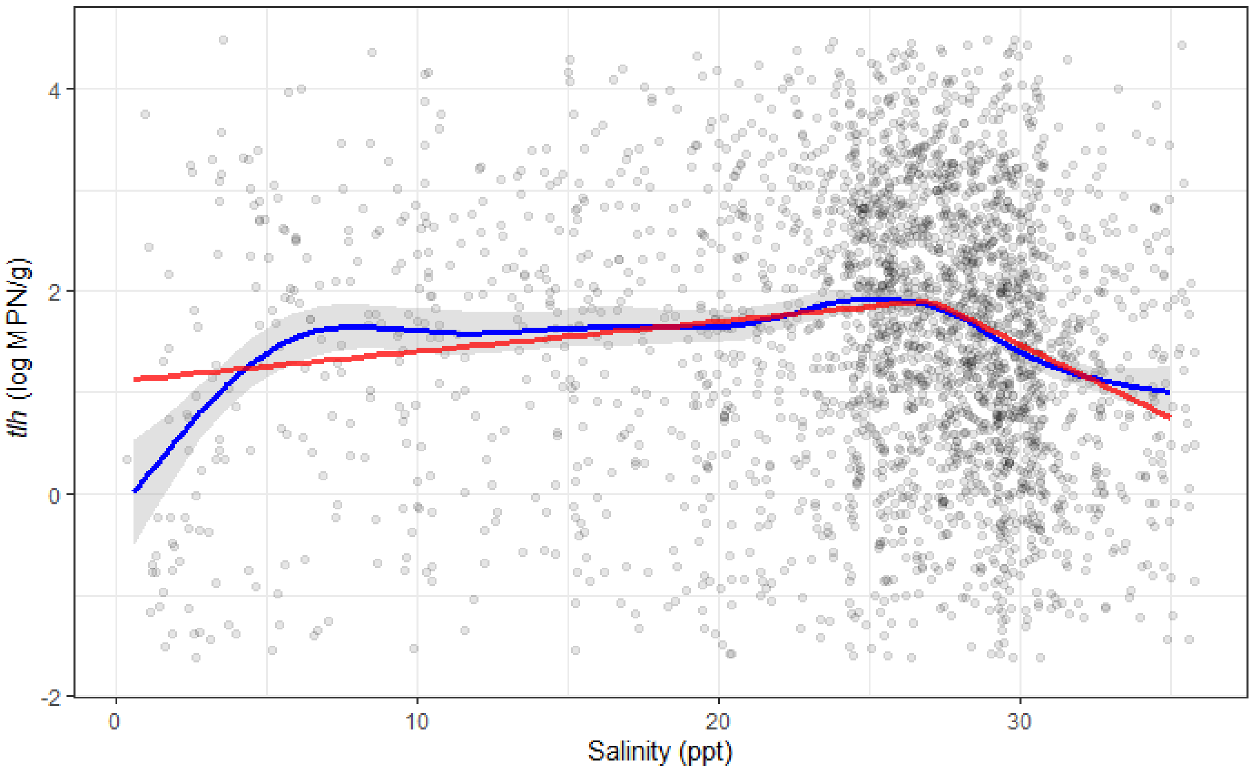


**Figure S7B.** Comparison of univariate regressions for salinity to *tlh* with linear regression with spline term (red) compared to smoothed conditional mean line of data (blue).

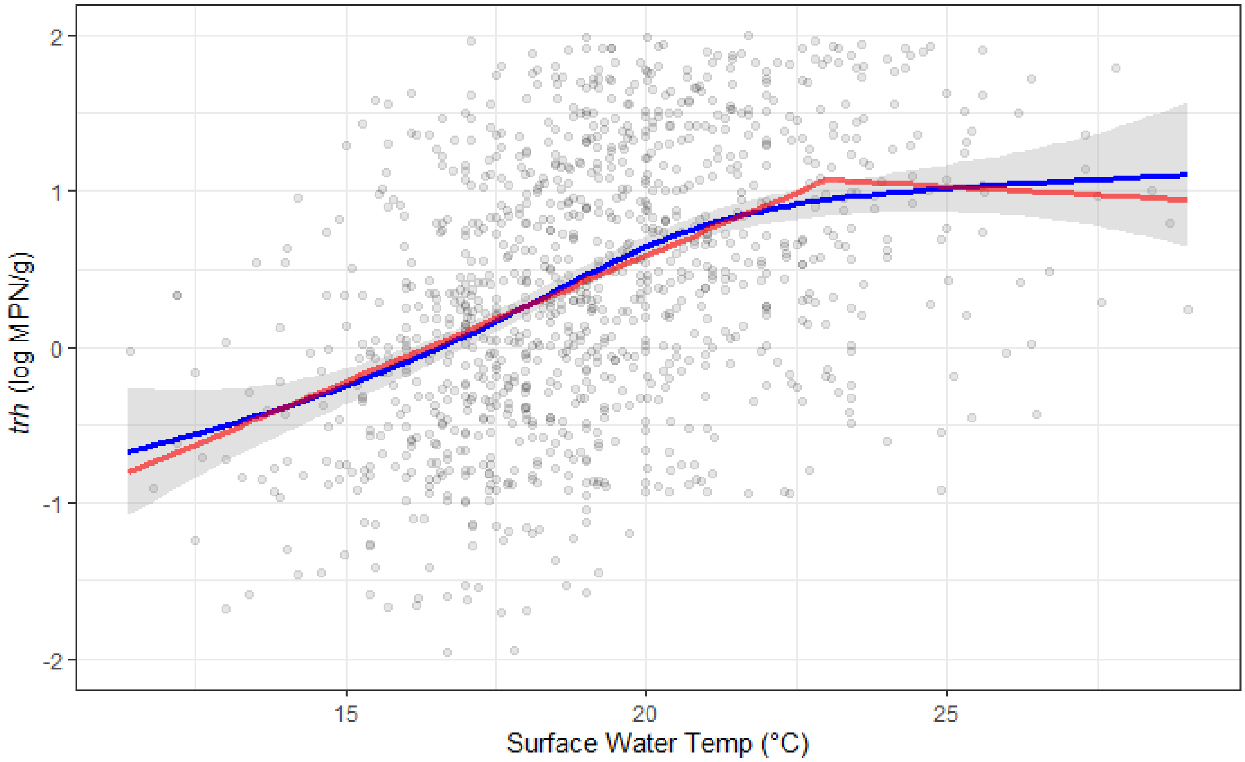


**Figure S7C.** Comparison of univariate regressions for surface water temperature to *trh* with linear regression with spline term (red) compared to smoothed conditional mean line of data (blue).

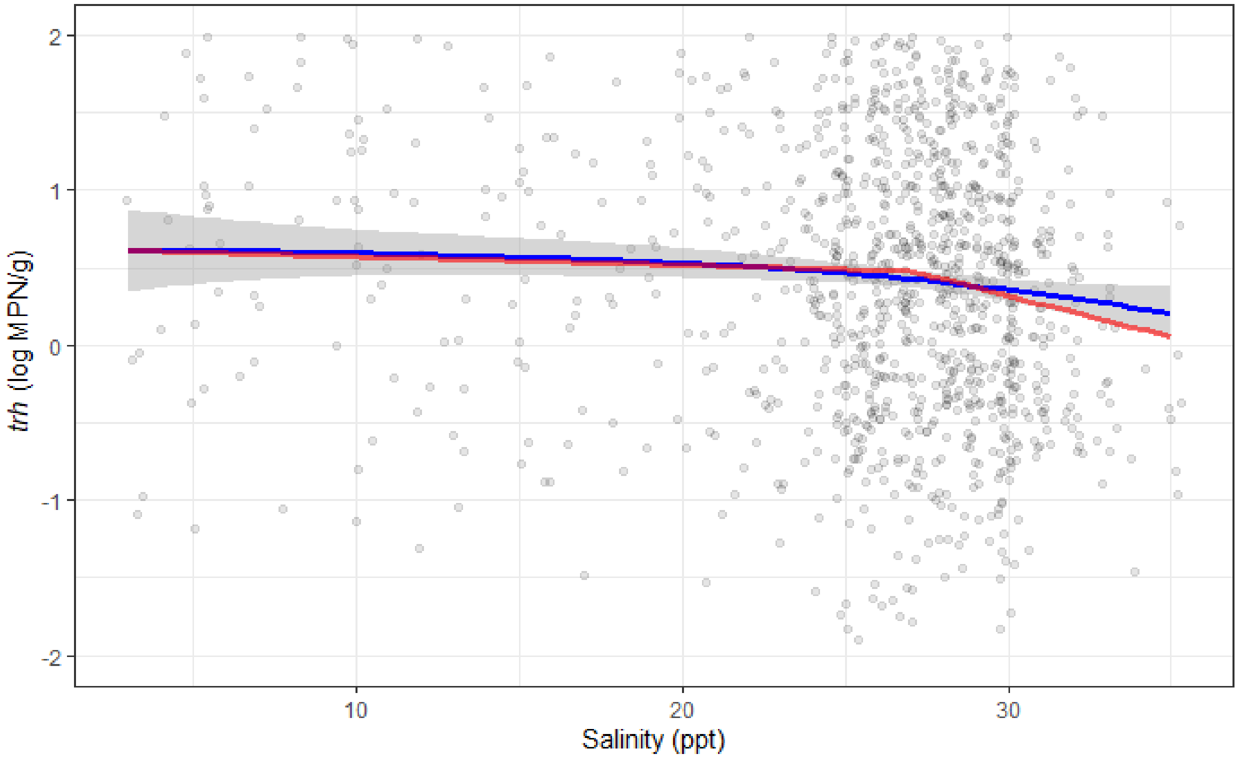


**Figure S7D.** Comparison of univariate regressions for salinity to *trh* with linear regression with spline term (red) compared to smoothed conditional mean line of data (blue).

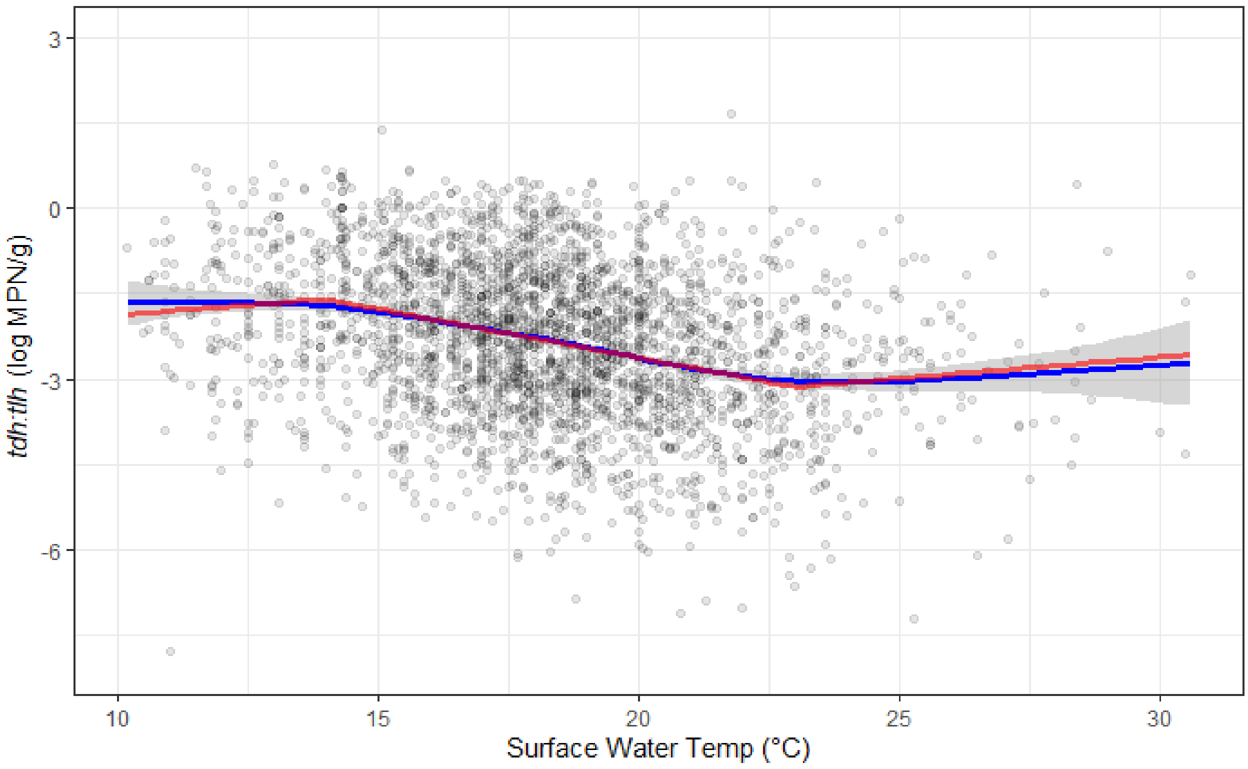


**Figure S7E.** Comparison of univariate regressions for salinity to *tdh*:*tlh* with linear regression with spline term (red) compared to smoothed conditional mean line of data (blue).

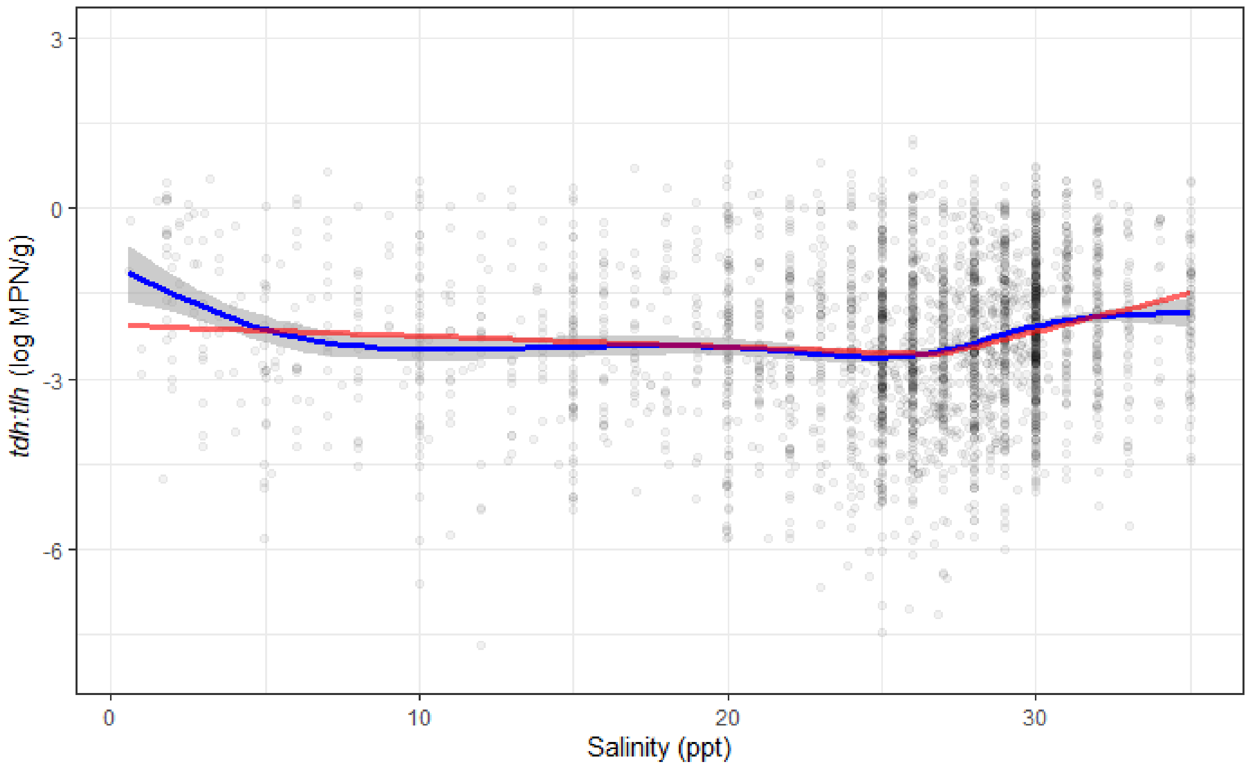


**Figure S7F.** Comparison of univariate regressions for salinity to *tdh*:*tlh* with linear regression with spline term (red) compared to smoothed conditional mean line of data (blue).

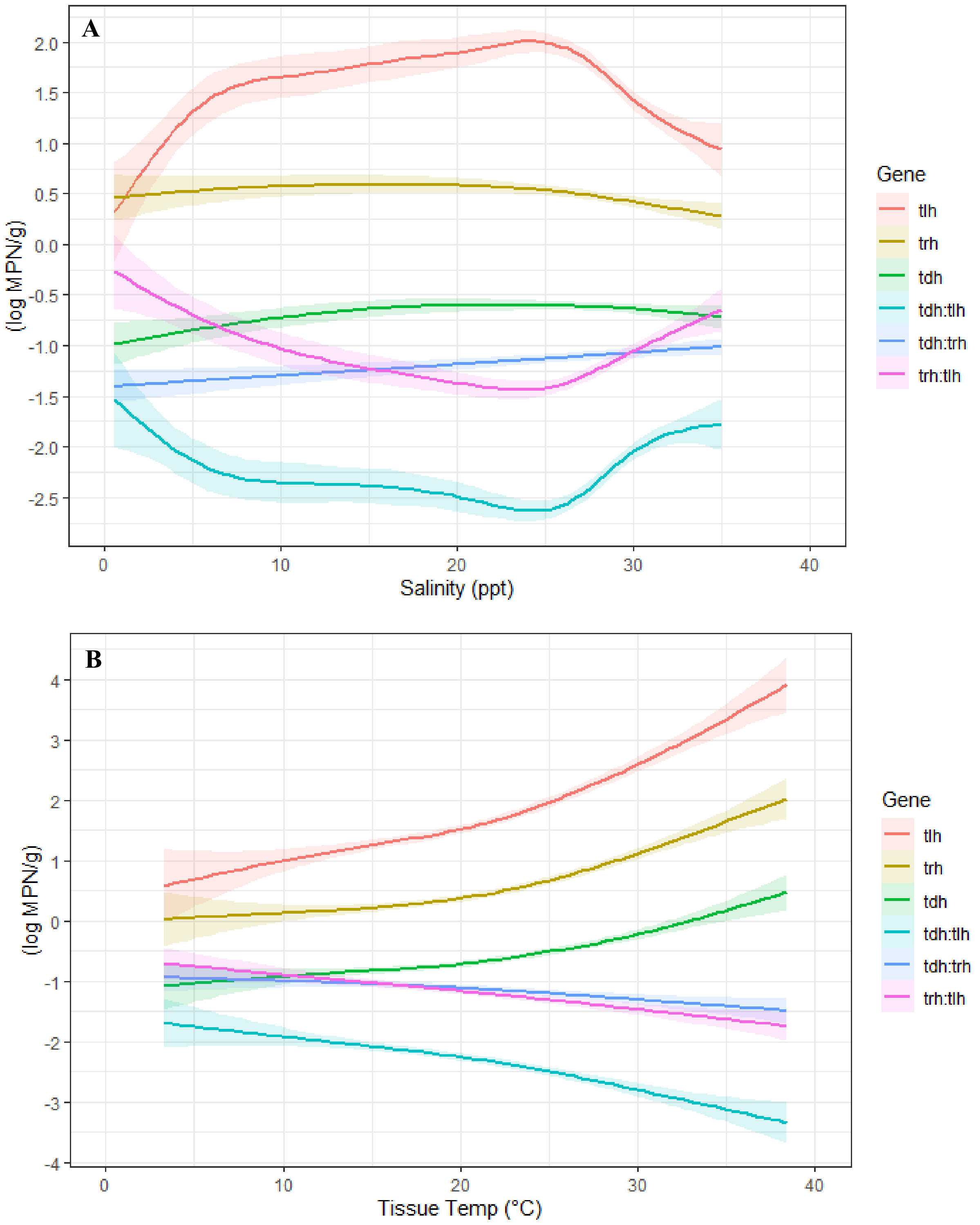


**Figure S8A.** Univariate lognormal regression analyses between a) salinity and b) tissue temperature, and genetic markers (representative imputation).

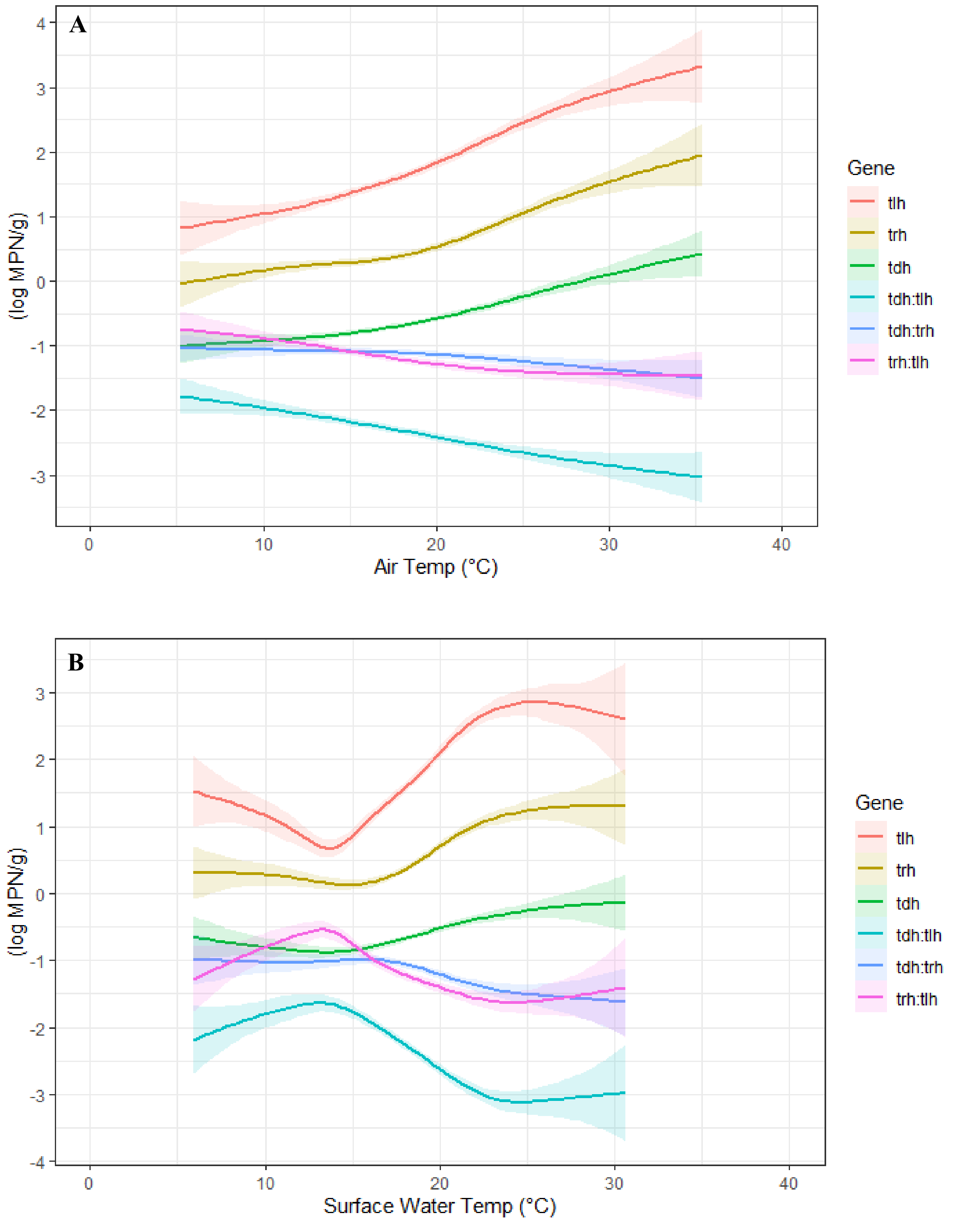


**Figure S8B.** Univariate lognormal regression analyses between a) ambient air temperature and b) surface water temperature, and genetic markers (representative imputation).

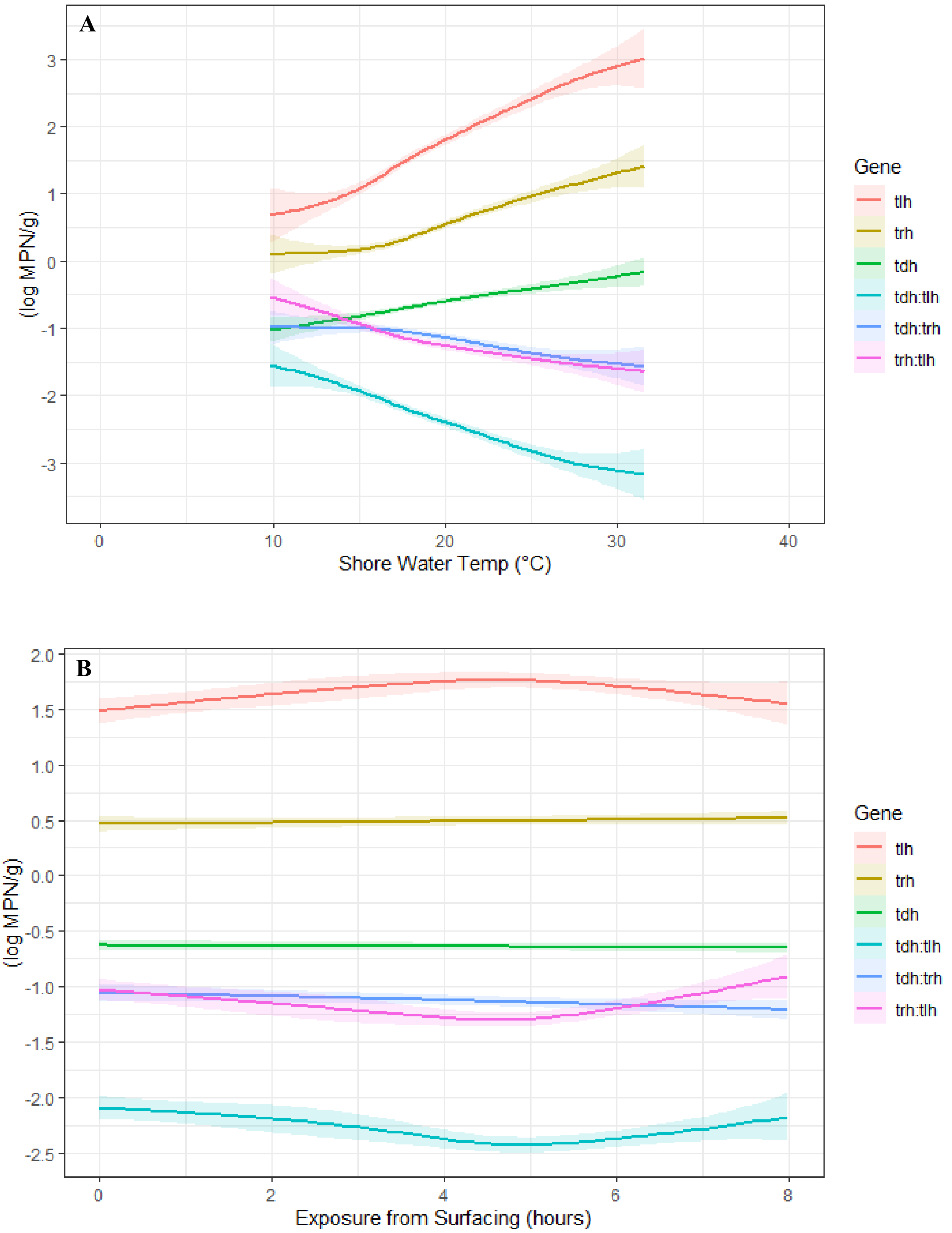


**Figure S8C.** Univariate lognormal regression analyses between a) shore water temperature and b) exposure from surfacing time, and genetic markers (representative imputation).

**Table S1.** Correlation matrices for time-indexed compared to lagged variables from 1-8 weeks lagged

Tissue Temperature

| | TTemp01 | TTemp02 | TTemp03 | TTemp04 | TTemp05 | TTemp06 | TTemp07 | TTemp08 | TissueTemp |
|---|---|---|---|---|---|---|---|---|---|
| TTemp01 | 1.00 | 0.18 | 0.28 | 0.09 | 0.20 | 0.05 | 0.10 | 0.04 | 0.18 |
| TTemp02 | 0.18 | 1.00 | 0.18 | 0.28 | 0.09 | 0.20 | 0.05 | 0.10 | 0.28 |
| TTemp03 | 0.28 | 0.18 | 1.00 | 0.18 | 0.28 | 0.09 | 0.20 | 0.05 | 0.09 |
| TTemp04 | 0.09 | 0.28 | 0.18 | 1.00 | 0.18 | 0.28 | 0.09 | 0.20 | 0.20 |
| TTemp05 | 0.20 | 0.09 | 0.28 | 0.18 | 1.00 | 0.18 | 0.28 | 0.09 | 0.05 |
| TTemp06 | 0.05 | 0.20 | 0.09 | 0.28 | 0.18 | 1.00 | 0.18 | 0.28 | 0.10 |
| TTemp07 | 0.10 | 0.05 | 0.20 | 0.09 | 0.28 | 0.18 | 1.00 | 0.18 | 0.04 |
| TTemp08 | 0.04 | 0.10 | 0.05 | 0.20 | 0.09 | 0.28 | 0.18 | 1.00 | 0.11 |
| TissueTemp | 0.18 | 0.28 | 0.09 | 0.20 | 0.05 | 0.10 | 0.04 | 0.11 | 1.00 |

Air Temperature

| | ATemp01 | ATemp02 | ATemp03 | ATemp04 | ATemp05 | ATemp06 | ATemp07 | ATemp08 | AirTemp |
|---|---|---|---|---|---|---|---|---|---|
| ATemp01 | 1.00 | 0.20 | 0.25 | 0.07 | 0.12 | -0.01 | 0.05 | 0.01 | 0.20 |
| ATemp02 | 0.20 | 1.00 | 0.20 | 0.25 | 0.07 | 0.12 | -0.01 | 0.05 | 0.25 |
| ATemp03 | 0.25 | 0.20 | 1.00 | 0.20 | 0.25 | 0.07 | 0.12 | -0.01 | 0.07 |
| ATemp04 | 0.07 | 0.25 | 0.20 | 1.00 | 0.20 | 0.25 | 0.07 | 0.12 | 0.12 |
| ATemp05 | 0.12 | 0.07 | 0.25 | 0.20 | 1.00 | 0.20 | 0.25 | 0.07 | -0.01 |
| ATemp06 | -0.01 | 0.12 | 0.07 | 0.25 | 0.20 | 1.00 | 0.20 | 0.25 | 0.05 |
| ATemp07 | 0.05 | -0.01 | 0.12 | 0.07 | 0.25 | 0.20 | 1.00 | 0.20 | 0.01 |
| ATemp08 | 0.01 | 0.05 | -0.01 | 0.12 | 0.07 | 0.25 | 0.20 | 1.00 | 0.07 |
| ATemp09 | 0.07 | 0.01 | 0.05 | -0.01 | 0.12 | 0.07 | 0.25 | 0.20 | 0.08 |
| ATemp10 | 0.08 | 0.07 | 0.01 | 0.05 | -0.01 | 0.12 | 0.07 | 0.25 | 0.13 |
| AirTemp | 0.20 | 0.25 | 0.07 | 0.12 | -0.01 | 0.05 | 0.01 | 0.07 | 1.00 |

Surface Water Temperature

| | SuTemp01 | SuTemp02 | SuTemp03 | SuTemp04 | SuTemp05 | SuTemp06 | SuTemp07 | SuTemp08 | SurfaceWaterTemp |
|---|---|---|---|---|---|---|---|---|---|
| SuTemp01 | 1.00 | 0.41 | 0.39 | 0.26 | 0.24 | 0.17 | 0.20 | 0.21 | 0.41 |
| SuTemp02 | 0.41 | 1.00 | 0.41 | 0.39 | 0.26 | 0.24 | 0.17 | 0.20 | 0.39 |
| SuTemp03 | 0.39 | 0.41 | 1.00 | 0.41 | 0.39 | 0.26 | 0.24 | 0.17 | 0.26 |
| SuTemp04 | 0.26 | 0.39 | 0.41 | 1.00 | 0.41 | 0.39 | 0.26 | 0.24 | 0.24 |
| SuTemp05 | 0.24 | 0.26 | 0.39 | 0.41 | 1.00 | 0.41 | 0.39 | 0.26 | 0.17 |
| SuTemp06 | 0.17 | 0.24 | 0.26 | 0.39 | 0.41 | 1.00 | 0.41 | 0.39 | 0.20 |
| SuTemp07 | 0.20 | 0.17 | 0.24 | 0.26 | 0.39 | 0.41 | 1.00 | 0.41 | 0.21 |
| SuTemp08 | 0.21 | 0.20 | 0.17 | 0.24 | 0.26 | 0.39 | 0.41 | 1.00 | 0.24 |
| SurfaceWaterTemp | 0.41 | 0.39 | 0.26 | 0.24 | 0.17 | 0.20 | 0.21 | 0.24 | 1.00 |

Salinity

| | Salt01 | Salt02 | Salt03 | Salt04 | Salt05 | Salt06 | Salt07 | Salt08 | Salinity |
|---|---|---|---|---|---|---|---|---|---|
| Salt01 | 1.00 | 0.50 | 0.47 | 0.43 | 0.47 | 0.44 | 0.46 | 0.47 | 0.50 |
| Salt02 | 0.50 | 1.00 | 0.50 | 0.47 | 0.43 | 0.47 | 0.44 | 0.46 | 0.47 |
| Salt03 | 0.47 | 0.50 | 1.00 | 0.50 | 0.47 | 0.43 | 0.47 | 0.44 | 0.43 |
| Salt04 | 0.43 | 0.47 | 0.50 | 1.00 | 0.50 | 0.47 | 0.43 | 0.47 | 0.47 |
| Salt05 | 0.47 | 0.43 | 0.47 | 0.50 | 1.00 | 0.50 | 0.47 | 0.43 | 0.44 |
| Salt06 | 0.44 | 0.47 | 0.43 | 0.47 | 0.50 | 1.00 | 0.50 | 0.47 | 0.46 |
| Salt07 | 0.46 | 0.44 | 0.47 | 0.43 | 0.47 | 0.50 | 1.00 | 0.50 | 0.47 |
| Salt08 | 0.47 | 0.46 | 0.44 | 0.47 | 0.43 | 0.47 | 0.50 | 1.00 | 0.50 |
| Salinity | 0.50 | 0.47 | 0.43 | 0.47 | 0.44 | 0.46 | 0.47 | 0.50 | 1.00 |

**Table S2.** Univariate and multivariate associations between *trh:tlh* and *tdh:trh* ratios and environmental covariates.

| | log *trh:tlh* | | log *tdh:trh* | |
|---|---|---|---|---|
| | **Univariate** | **Multivariate** | **Univariate** | **Multivariate** |
| **Ecological characteristic** | | | | |
| Salinity (ppt) - 3 week lag | - | - | 0.02 (0.01, 0.04) | 0.01 (0.00, 0.03) |
| *1 ppt - 27 ppt* | -0.20 (-0.48, 0.07) | -0.12 (-0.40, 0.15) | - | - |
| *27 ppt - 35 ppt* | 0.39 (0.05, 0.73) | 0.39 (0.06, 0.72) | - | - |
| Tissue Temp (°C) | 0.00 (0.00, 0.01) | * | -0.03 (-0.04, -0.02) | * |
| Air Temp (°C) - 1 week lag | -0.01 (-0.02, 0.00) | * | -0.02 (-0.05, -0.02) | * |
| Surface Water Temp (°C) | -0.02 (-0.03, 0.00) | -0.02 (-0.04, 0.00) | -0.08 (-0.10, -0.06) | -0.06 (-0.09, -0.04) |

*Results are displayed as the log-transformed, pooled parameter estimates of the model with associated 95% confidence intervals. Reported associations are adjusted for region and year. * Indicated null effect (0) and exclusion from multivariate model.*

**Table S3.** Model Random Effects for Zone and Sample Site Year Group (SSYG)

| | log *trh:tlh* | log *tdh:trh* |
|---|---|---|
| **Random Effects** | | |
| **Univariate** | | |
| Salinity (ppt) | | |
| *Random Intercept - Zone* | 0.33 | 0.01 |
| *Random Intercept - SSYG* | 0.23 | 0.23 |
| Tissue Temp (°C) | | |
| *Random Intercept - Zone* | 0.37 | 0.00 |
| *Random Intercept - SSYG* | 0.25 | 0.25 |
| Air Temp (°C) | | |
| *Random Intercept - Zone* | 0.36 | 0.00 |
| *Random Intercept - SSYG* | 0.25 | 0.25 |
| Surface Water Temp (°C) | | |
| *Random Intercept - Zone* | 0.35 | 0.00 |
| *Random Intercept - SSYG* | 0.24 | 0.25 |
| **Multivariate** | | |
| *Random Intercept - Zone* | 0.30 | 0.00 |
| *Random Intercept - SSYG* | 0.11 | 0.01 |

*Estimate of random intercept effect size for univariate and multivariate models.*

**Figure S9.** ACF and PACF of model residuals before and after applying nested ARMA structure to models

*tlh* model residual correlation **before** ARMA

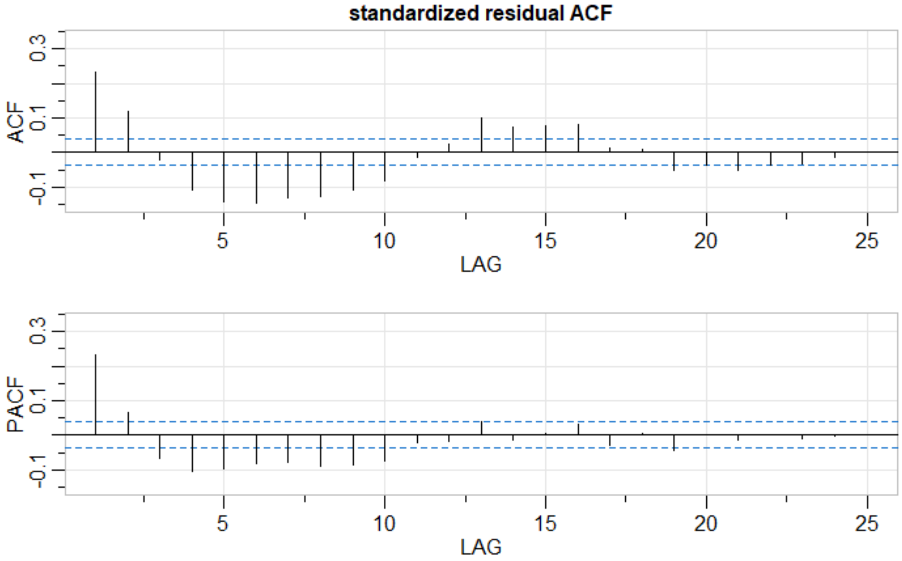


*tlh* model residual correlation **after** ARMA structure

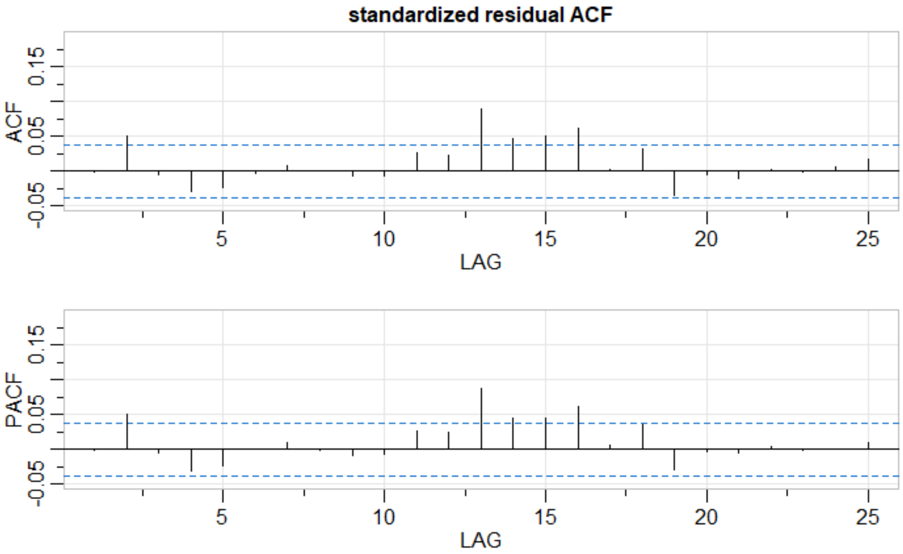

*trh* model residual correlation **before** ARMA structure

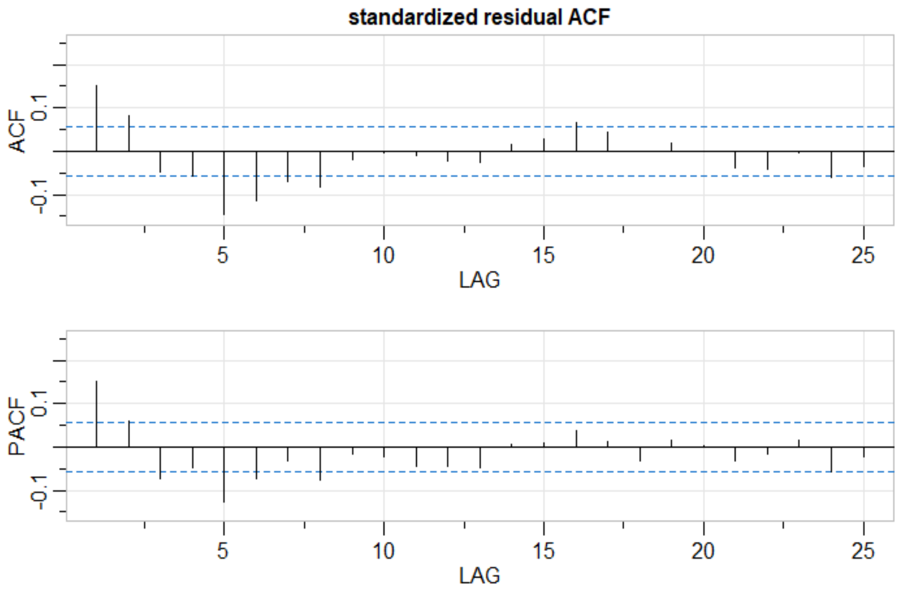


*trh* model residual correlation **after** ARMA structure

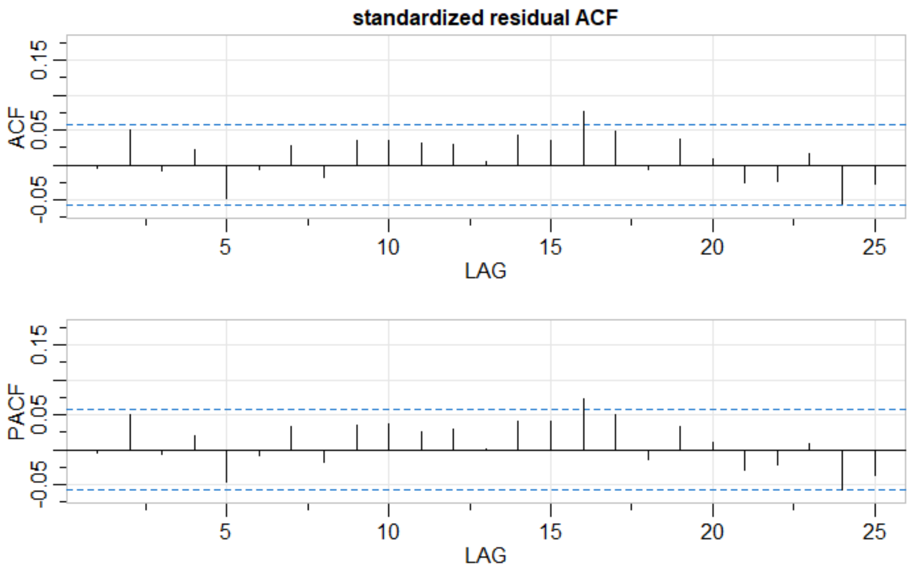

*tdh* model residual correlation **before** ARMA structure

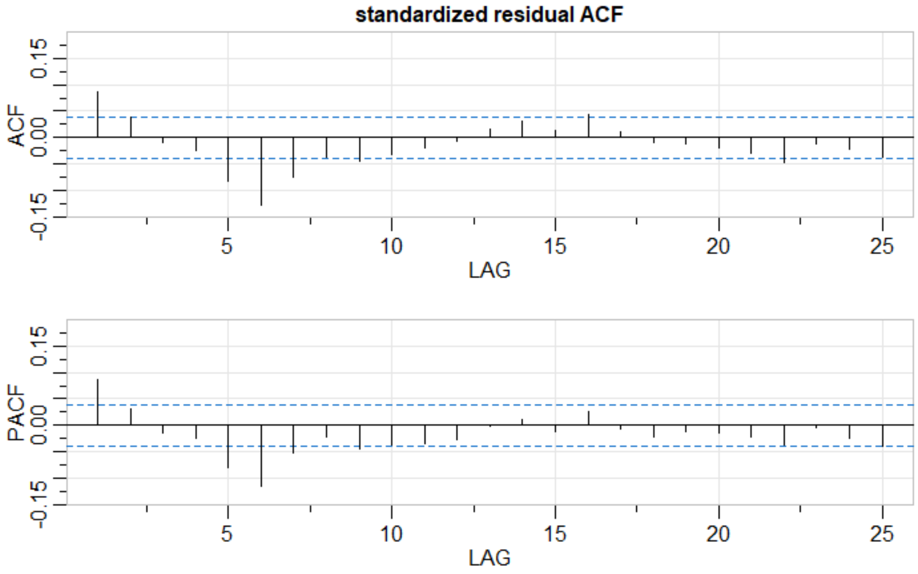


*tdh* model residual correlation **after** ARMA structure

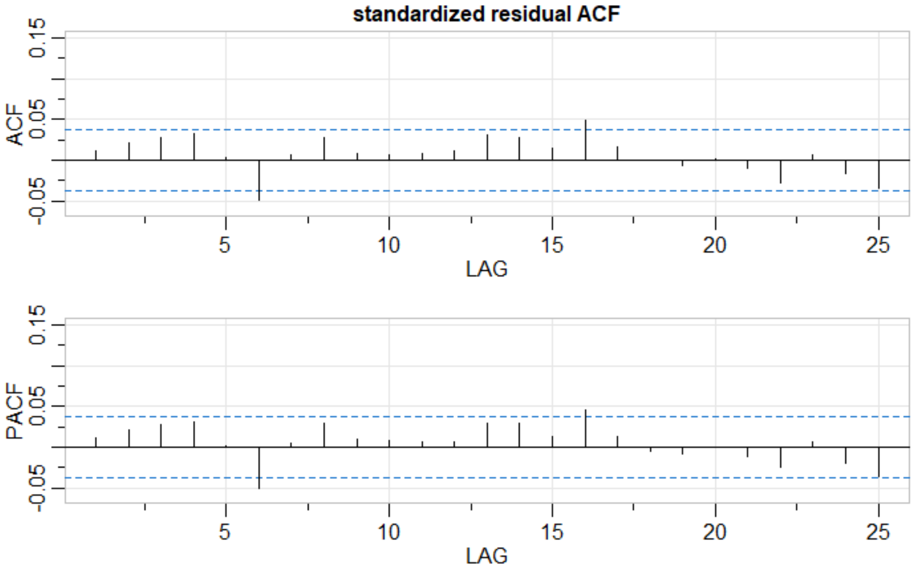

*tdh*:*tlh* model residual correlation **before** ARMA structure

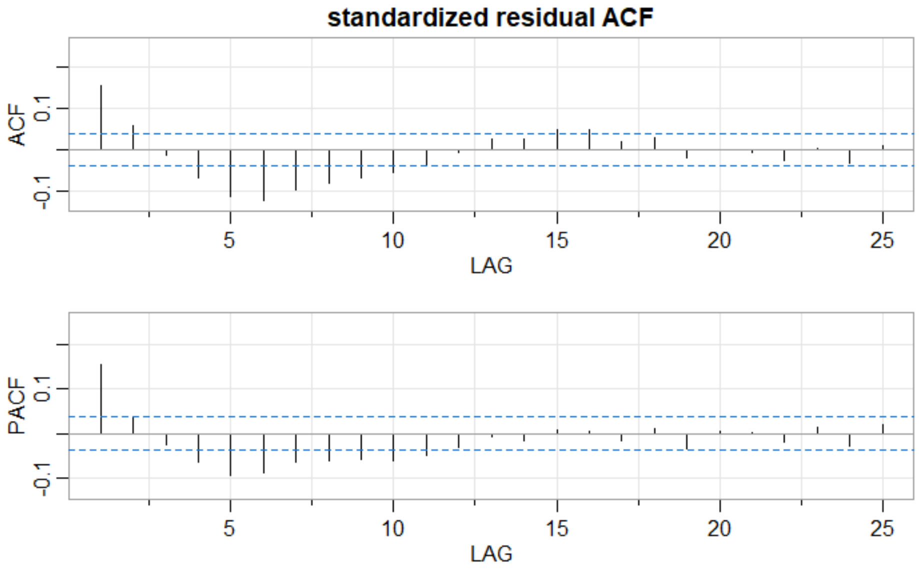


*tdh*:*tlh* model residual correlation **after** ARMA structure

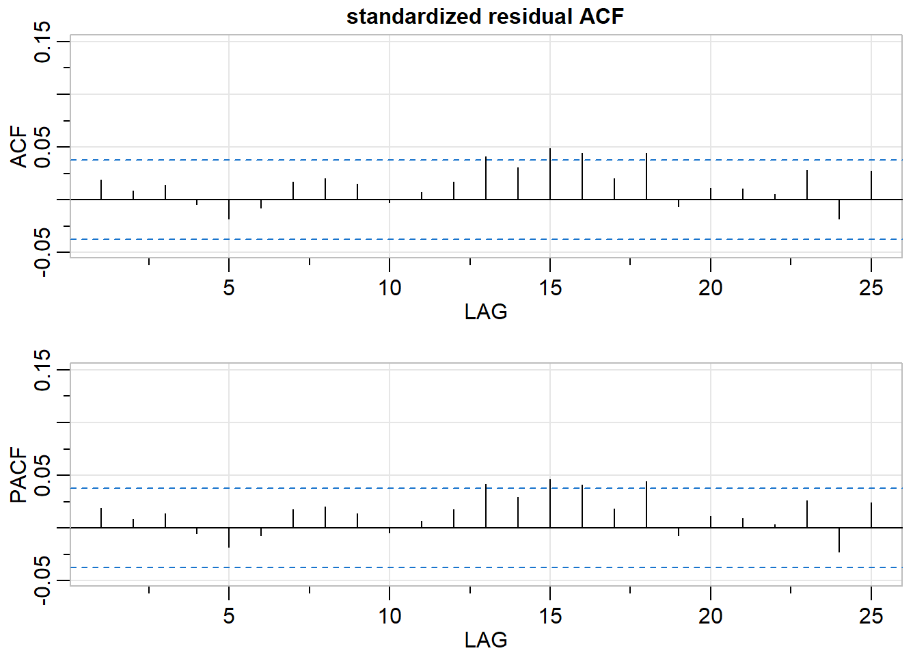

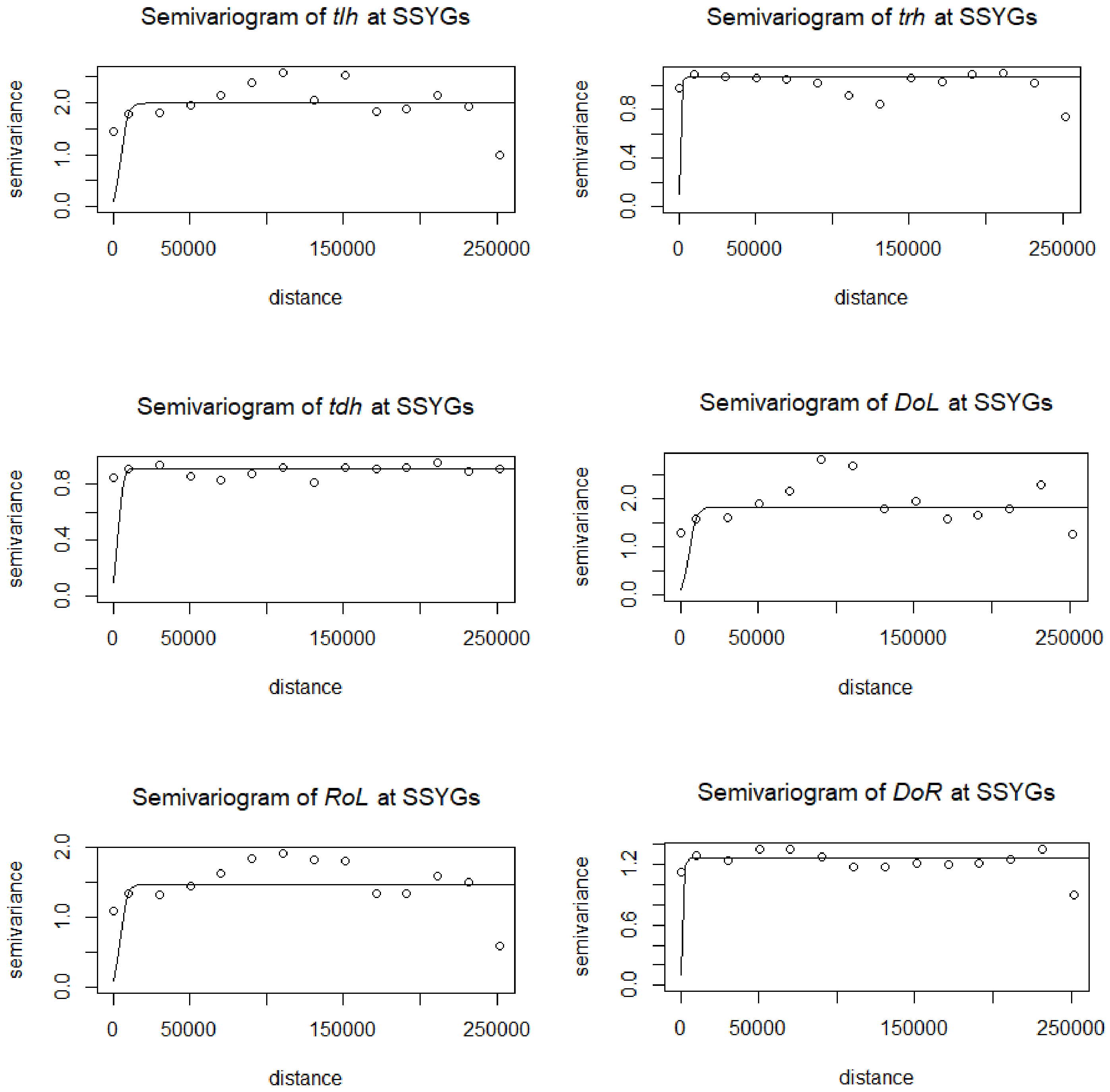


Figure S10. Variograms of spatial autocorrelation of environmental characteristics and genetic markers from model using non-Euclidian water distance function.
